# Pharmacologically inducing regenerative cardiac cells by small molecule drugs

**DOI:** 10.1101/2023.10.24.563872

**Authors:** Wei Zhou, Kezhang He, Chiyin Wang, Pengqi Wang, Dan Wang, Bowen Wang, Han Geng, Hong Lian, Tianhua Ma, Yu Nie, Sheng Ding

## Abstract

Adult mammals, unlike some lower organisms, lack the ability to regenerate damaged hearts through cardiomyocytes (CMs) dedifferentiation into cells with regenerative capacity. Developing conditions to induce such naturally unavailable cells with potential to proliferate and differentiate into CMs, i.e., regenerative cardiac cells (RCCs), in mammals will provide new insights and tools for heart regeneration research. In this study, we demonstrate that a two-compound combination, CHIR99021 and A-485 (2C), effectively induces RCCs from human embryonic stem cell (hESC)-derived TNNT2^+^ CMs *in vitro*, as evidenced by lineage tracing experiments. Functional analysis shows that these RCCs express a broad spectrum of cardiogenesis genes and have the potential to differentiate into functional CMs, endothelial cells (ECs), and smooth muscle cells (SMCs). Importantly, similar results were observed in neonatal rat CMs both *in vitro* and *in vivo*. Remarkably, administering 2C in adult mouse hearts significantly enhances survival and improves heart function post-myocardial infarction. Mechanistically, CHIR99021 is crucial for the transcriptional and epigenetic activation of genes essential for RCC development, while A-485 primarily suppresses H3K27Ac and particularly H3K9Ac in CMs. Their synergistic effect enhances these modifications on RCC genes, facilitating the transition from CMs to RCCs. Therefore, our findings demonstrate the feasibility and reveal the mechanisms of pharmacological induction of RCCs from endogenous CMs, which could offer a promising regenerative strategy to repair injured hearts.

---

Lower organisms like zebrafish exhibit a remarkable capacity for cardiac regeneration through CM dedifferentiation, characterized by the reactivation of embryonic cardiogenic genes and disassembly of sarcomeric structures (Kikuchi & Poss, 2010; Lepilina et al., 2006). In contrast, mammals show limited cardiac regeneration, typically restricted to the early postnatal period (PN1 to PN6) in mice, beyond which the regenerative ability sharply declines (Porrello et al., 2011). This stark difference underscores the need for novel strategies to induce a regenerative state in adult mammalian CMs, akin to that observed in zebrafish, to overcome the inherent barriers to CM dedifferentiation.

During mammalian heart development, CMs arise predominantly from the first heart field (FHF) and the second heart field (SHF) (Srivastava, 2006), with SHF cells marked by the pioneer transcription factor ISL1 proliferating and differentiating into major cardiovascular cell types, CMs, SMCs and ECs during embryonic development (Bu et al., 2009; Moretti et al., 2006). Notably, ISL1^+^ cells in neonatal mouse hearts show potential for expansion and differentiation into CMs under conducive *in vitro* conditions (Laugwitz, 2005). Given that ISL1 transcriptionally governs the expression of a suite of cardiac genes, such as *NKX2-5* (Ma et al., 2008), *FGF10* (Watanabe et al., 2012), which are essential for embryonic cardiogenesis, inducing ISL1 expression in CMs could be a critical determinant in the induction of cells possessing regenerative capacity.

Aiming to replicate this regenerative capacity, our study focused on identifying small molecules capable of inducing ISL1 expression in CMs. Through a combinatorial screening approach, we discovered a novel combination of CHIR99021 and A-485, or 2C, that efficiently and unprecedentedly induces dedifferentiation of human CMs into RCCs. These RCCs exhibited disassembled sarcomeric structures, high expression of embryonic cardiogenic genes, and an increased number of CMs through re-differentiation *in vitro*. Further investigations showed that 2C robustly generates RCCs in adult mouse hearts and improves cardiac function in mice experiencing myocardial infarction. This proof-of-concept discovery demonstrates that a simple combination of small molecule drugs can endow the mammalian heart with regenerative capacity by reprogramming CMs into RCCs.

## Efficient induction of dedifferentiation in hESC-derived CMs by 2C treatment

To obtain an adequate number of CMs for small molecule screening, we differentiated hESCs into CMs (Movie 1, Figure 1—figure supplement 1A) following a well-established step-wise protocol (Lian et al., 2013). Nearly homogeneous contracting CMs were observed on day 10 of differentiation, consistent with previous reports. After purification with glucose-depleted medium containing abundant lactate (Figure 1— figure supplement 1B), highly pure TNNT2^+^ CMs were obtained. These were subsequently dissociated and seeded into 96-well plates. Once the CMs resumed contraction, they were treated with individual small molecules from a collection of over 4,000 compounds for 3 days (Figure 1—figure supplement 1C), and then fixed and immunostained for ISL1. Using a high-content imaging (HCI) and analysis system, five compounds were initially identified as potential inducers of dedifferentiation in CMs, indicated by induced sarcomere disassembly and ISL1 expression (Figure 1—figure supplement 1D). Further comparison revealed that the unique combination of CHIR99021 and A-485 (2C) most efficiently induced ISL1 expression in TNNT2^+^ CMs (Figure 1—figure supplement 1E). Notably, while A-485 was screened at a concentration of 10 µM, its optimal working concentration in subsequent experiments was determined to be 0.5 µM based on titration experiments (Figure 1—figure supplement 1F-G). Of note, compound I-BET-762 also showed some capacity to induce ISL1 expression. However, it was less effective than A-485 in combination with CHIR99021 (Figure 1—figure supplement 1H).

**Figure 1.**
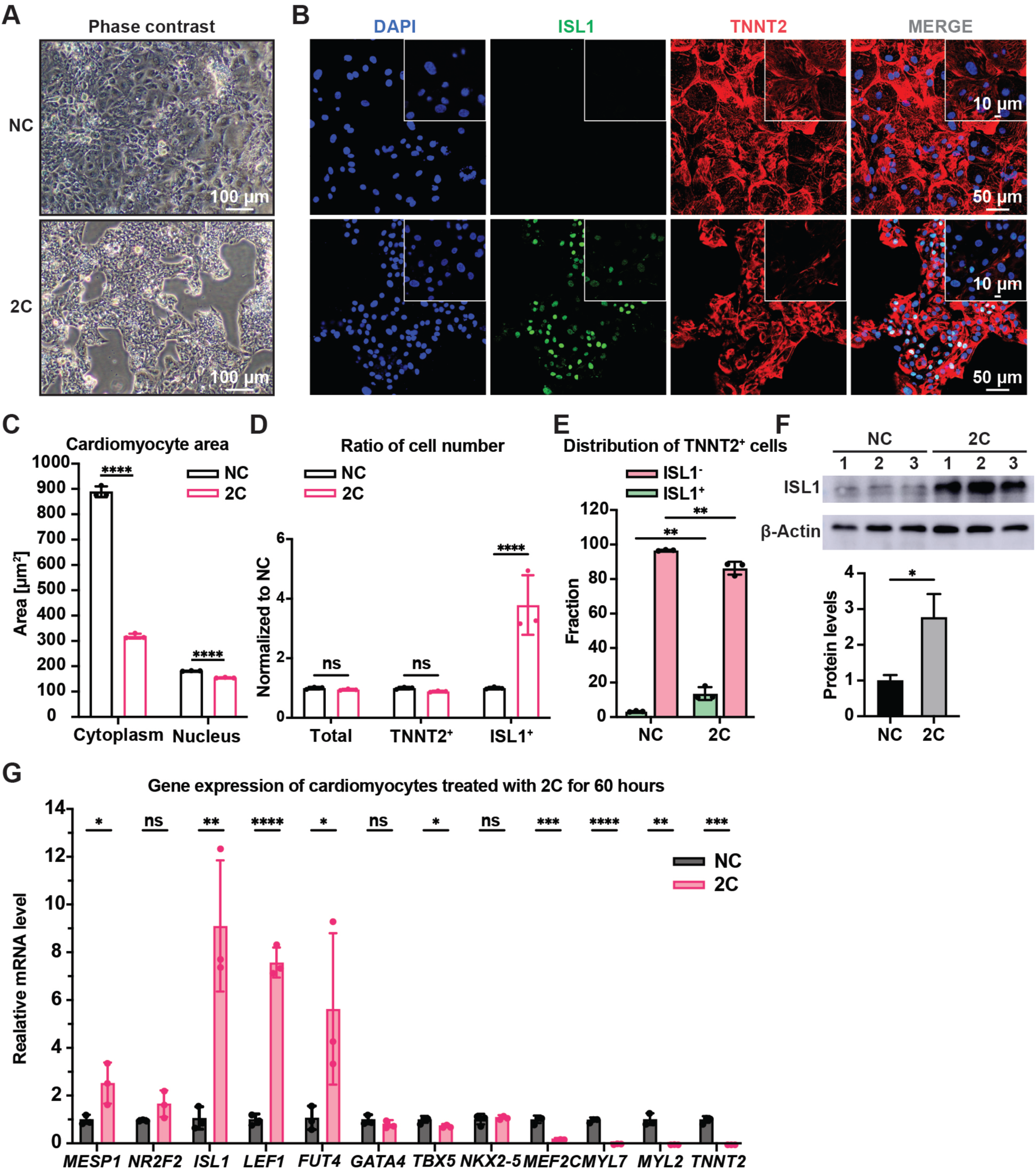
2C treatment induced dedifferentiation of hESC-derived CMs toward ISL1-expressing cells. Cells induced from hESC-derived CMs by treatment with DMSO (NC) or 2C for 60 hours. **(A)** Phase contrast images showing cell morphology. **(B)** Immunofluorescence staining of the ISL1 (ISL1, green), and the CM marker cardiac troponin T (TNNT2, red) in the cells. Nuclei were stained by DAPI (4′,6-diamidino-2-phenylindole) and presented in DNA blue. **(C)**, Cytosolic and nuclear areas of the cells. Data are shown as mean ± SD. (n=3 independent experiments, represented as dots). Two-way ANOVA with Dunnett’s multiple comparisons test. ****P < 0.0001. **(D)** Changes in cell number after 2C treatment. Total cell number, TNNT2^+^ cell number and ISL1^+^ cell number were normalized to the negative control DMSO (NC). Data are shown as mean ± SD (n=3 independent experiments, represented as dots). Two-way ANOVA with Šidák’s multiple comparisons test. ns, not significant (P > 0.05), ****P < 0.0001. **(E)** Fraction of ISL1^+^ cells in TNNT2^+^ cells. Data are shown as mean ± SD (n=3 independent experiments, represented as dots). Two-way ANOVA with Šidák’s multiple comparisons test. **P < 0.01. **(F)** Western blot and quantitative analysis of ISL1 expression in DMSO (NC) or 2C-treated CMs for 60 hours. Data are shown as mean ± SD. Unpaired t test. *P < 0.05. **(G)** Relative gene expression of embryonic cardiogenesis marker genes (*MESP1*, *ISL1*, *NR2F2*, *FUT4*, and *LEF1*), pan-cardiac genes (*GATA4*, *TBX5*, and *NXK2-5*), and CM marker genes (*MEF2C*, *TNNT2*, *MYL2*, and *MYL7*) in the cells treated by DMSO (NC) or 2C for 60 hours (60h). Data are shown as mean ± SD (n=3 independent experiments, represented as dots). Multiple unpaired t tests. ns, not significant (P > 0.05), *P < 0.05, **P < 0.01, ***P < 0.001, ****P < 0.0001.

When CMs with well-organized sarcomeres were treated with 2C *in vitro*, cells started to exhibit a dedifferentiation-associated phenotype, such as reduction of cell size after 24 hours, growing as clusters after 48 hours, and forming colonies after 60 hours (Figure 1A and Figure 1—figure supplement 2A). During this period, the expression of TNNT2 and MYL2 gradually downregulated while ISL1 expression and the percentage of ISL1^+^ cells increased (Figure 1—figure supplement 2B). Compared to untreated CMs, significant sarcomere disassembly and reduction of cytoplasmic/nuclear area were observed in cells treated with 2C for 60 hours (Figure 1B-C). In parallel, the number and percentage of TNNT2^+^ cells remained constant while ISL1^+^ cells increased remarkably (Figure 1D-E and Figure 1—figure supplement 2C), accompanied by a ∼3-fold increase in ISL1 expression at both mRNA and protein levels (Figure 1F and Figure 1—figure supplement 2B). Other early embryonic cardiogenesis genes, including MESP1, LEF1 (Klaus et al., 2007), FUT4 (Wang et al., 2019), and NR2F2 (Churko et al., 2018), were also highly expressed following 2C treatment (Figure 1G and Figure 1—figure supplement 2D). Furthermore, 2C treatment significantly decreased the expression of CM-specific genes, such as TNNT2, MYL2, MYL7, and MEF2C, while pan-cardiac transcription factors GATA4, TBX5, and NKX2-5 showed less effect (Figure 1G and Figure 1—figure supplement 2D). Notably, 2C induced ISL1 expression even in mature CMs derived from hESCs treated with ZLN005 (Liu et al., 2020) (Figure 1—figure supplement 2E). These results indicate that 2C treatment effectively reprograms CMs towards a dedifferentiated, embryonic-like state.

## 2C induced cells possess regenerative capacity

To confirm the regenerative capacity of 2C-induced cardiac cells, we investigated cell proliferation using a BrdU incorporation assay. Immunostaining showed the co-localization of ISL1 and BrdU labeling in these cells (Figure 2A). Statistical analysis revealed a significant decrease in the nuclear area and an increase in the number of ISL1^+^/BrdU^+^ positive cells following 2C treatment (Figure 2B-C). We then examined the potential for re-differentiation of 2C-induced cardiac cells into CMs. As expected, upon withdrawal of 2C and subsequent culture in CM media for 3 days (60h+3d), we observed the emergence of spontaneously contracting cells exhibiting CM-specific morphology and typical cytoplasmic/nuclear area size (Movie 2 and Figure 2D-E). These cells downregulated early embryonic cardiogenesis genes and upregulated CM-specific genes, with no noticeable changes in pan-cardiac genes (Figure 2—figure supplement 1A). Remarkably, there was a notable increase (roughly 1.4-fold) in the number of re-differentiated CMs with clear TNNT2 staining relative to the initial CMs before 2C treatment (Figure 2F), demonstrating the regenerative potential in 2C-induced cardiac cells. Additionally, 2C-induced cardiac cells could differentiate into SMA^+^ SMCs or CD31^+^/VE-Cadherin^+^ ECs in the presence of PDGF-BB and TGF-β1 or VEGF, bFGF and BMP4, respectively (Figure 2G-H and Figure 2—figure supplement 1B). In contrast, these genes expressed were not detectable in DMSO-treated cells. Collectively, our findings indicate that 2C treatment facilitates the conversion of CMs into cardiac cells with regenerative capability, including proliferative potential and differentiation into the three major cardiovascular cell types, thus named regenerative cardiac cells (RCCs).

**Figure 2.**
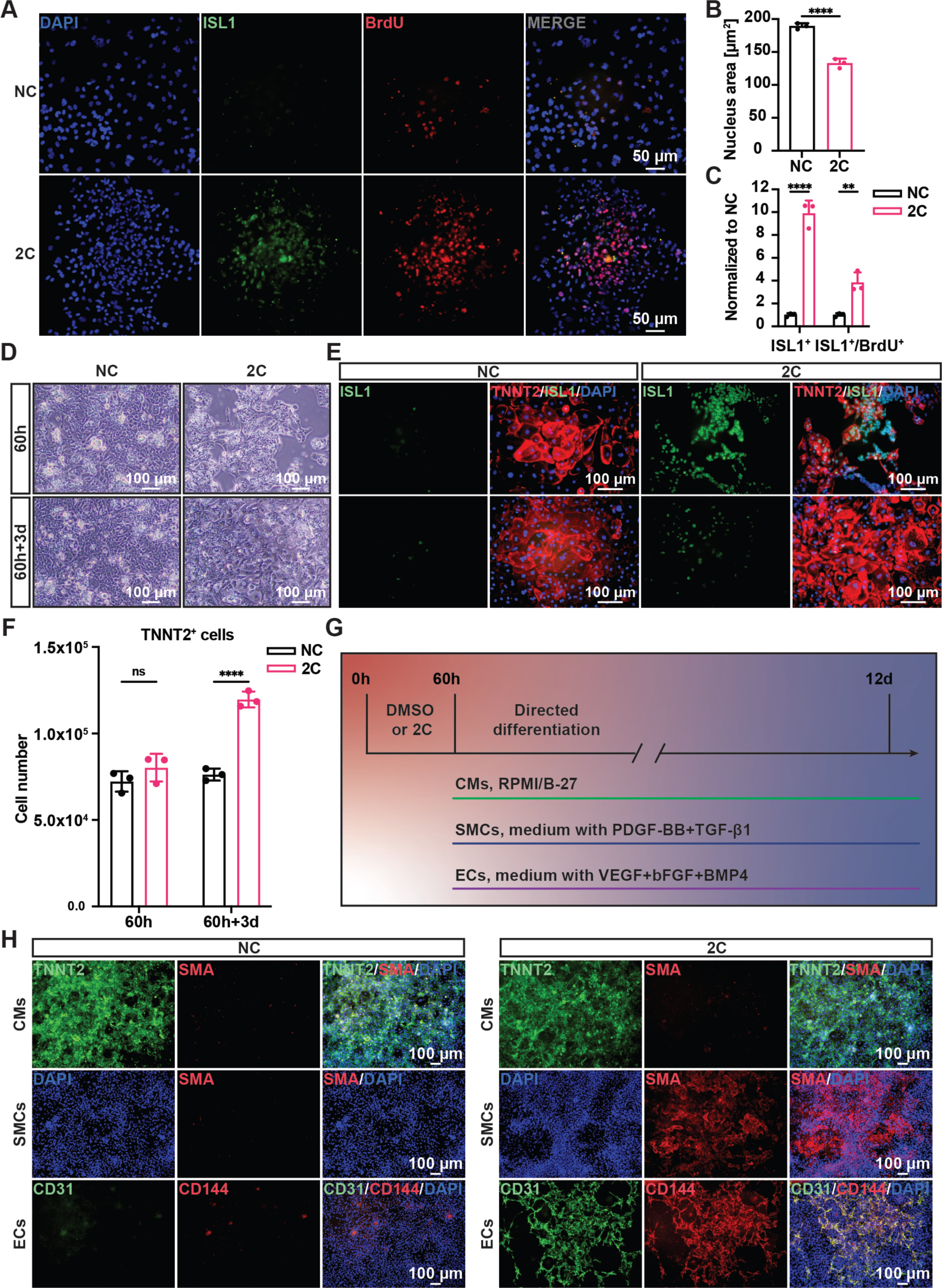
Regenerative ability of 2C-induced RCCs. **(A-C)** Immunofluorescence staining (**A**) and statistical analysis (**B** and **C**) of the ISL1 (green) and BrdU (red) double positive RCCs induced from CMs by treatment with DMSO (NC) or 2C for 60 hours. The ISL1^+^ cell number and ISL1^+^/BrdU^+^ cell number were normalized to the negative control DMSO (NC). DAPI (4′,6-diamidino-2-phenylindole) staining labeled nuclei as blue. Data are shown as mean ± SD (n=3 independent experiments, represented as dots). Multiple unpaired t tests in (**B**), ****P < 0.0001. Two-way ANOVA with Šidák’s multiple comparisons test in (**C**), **P < 0.01, ****P < 0.0001. **(D)** Phase contrast images of hESC-derived CMs treated by DMSO (NC) or 2C for 60 hours (60h) and subsequently cultured in the absence of 2C for another 3 days (60h+3d). **(E)** Immunostaining showed the expression of ISL1 (green) and TNNT2 (red) in the cells under the same condition in (**D**). **(F)** Statistical analysis of TNNT2^+^ cell numbers under the same condition in (**D**). Data are shown as mean ± SD (n=3 independent experiments, represented as dots). Two-way ANOVA with Šidák’s multiple comparisons test. ns, not significant (P > 0.05), ****P < 0.0001. **(G)** Schematic diagram of directed differentiation of 2C-induced RCCs towards cardiomyocytes (CMs), smooth muscle cells (SMCs) and endothelial cells (ECs). **(H)** Immunostaining showed the expression of EC markers (CD31, green and CD144, red), SMC marker (SMA, red), and CM marker (TNNT2, green). DAPI (4′,6-diamidino-2-phenylindole) staining labeled nuclei as blue.

## Lineage tracing demonstrated that 2C induced RCCs dedifferentiated from TNNT2^+^ CMs

Although TNNT2^+^ CMs purified using a lactate-based culture medium were nearly homogeneous populations, a small fraction (<4%) of purified cells still expressing ISL1 (Figure 1—figure supplement 1B). To ascertain that the RCCs induced by 2C treatment were indeed dedifferentiated from TNNT2^+^ CMs rather than originating from the proliferation of residual ISL1^+^ cells, we conducted lineage-tracing experiments. Using an ISL1^mCherry/+^ H9 hESC (K9) line established through CRISPR-based knock-in (Figure 3—figure supplement 1A-D), we verified that mCherry expression accurately mirrored endogenous ISL1 expression (Figure 3A-C). Following the purification of K9-derived CMs and the selection of mCherry-negative cells by FACS (Figure 3D-E), these cells were confirmed to be TNNT2^+^ CMs (Figure 3—figure supplement 1E). Upon treatment of these mCherry-negative CMs with 2C for 60 hours, we observed remarkable morphological changes in mCherry^+^/ISL1^+^ RCCs compared to mCherry^−^/TNNT2^+^ CMs (Figure 3F), accompanied by a dramatic downregulation of TNNT2 and MYL2 expression and a notable upregulation of ISL1 and mCherry expression (Figure 3G). FACS analysis further validated that 2C could convert K9-derived mCherry-negative CMs into mCherry^+^/ISL1^+^ cells (Figure 3H-I). Similar findings were obtained using the HUES7 hESC line with an ISL1-mCherry knock-in reporter (K7) (Figure 3—figure supplement 2), robustly demonstrating that 2C induces ISL1 expression in TNNT2^+^ CMs, effectively generating RCCs.

**Figure 3.**
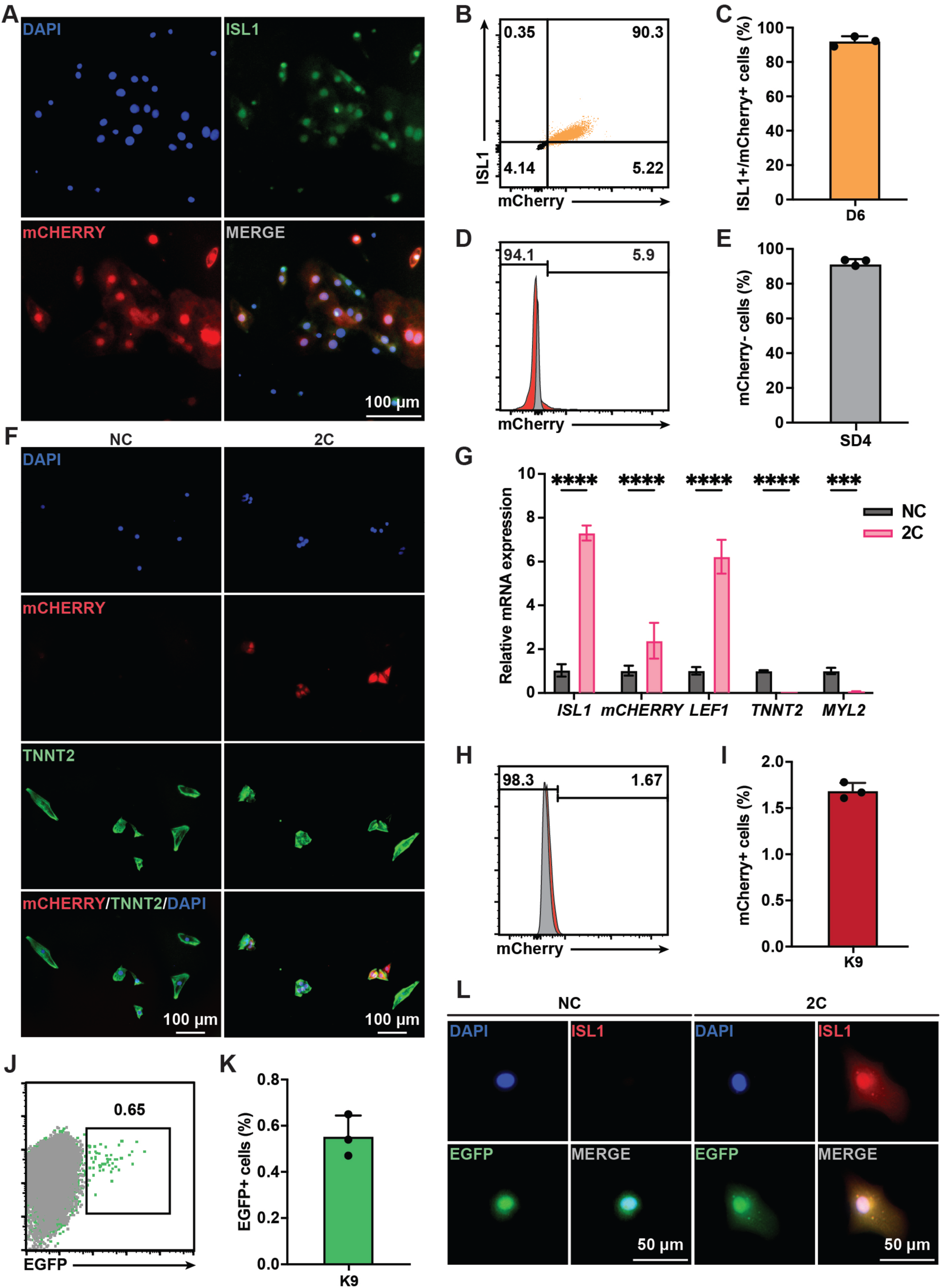
Lineage tracing demonstrated 2C induced dedifferentiation of TNNT2^+^ CMs to ISL1-expressing RCCs. **(A)** Immunofluorescence images showing expression of endogenous ISL1 (green) and ISL1-mCherry (red) reporter in the cells differentiated from K9 hESC KI reporter line at day 6 (D6). DAPI (4′,6-diamidino-2-phenylindole) staining labeled nuclei as blue. **(B-C)** Flow cytometry analysis of the percentage of mCherry^+^/ISL1^+^ cells in the cells differentiated from K9 at D6. **(D-E)** Flow cytometry analysis of the percentage of mCherry-negative cells at selection day 4 (SD4) in lactate purification medium. **(F)** Cells induced from mCherry-negative CMs by treatment with or without 2C for 60 hours. Images showing the expression of mCHERRY (red) and TNNT2 (green) in the cells. **(G)** Relative gene expression of *ISL1*, *mCHERRY*, *LEF1*, *TNNT2* and *MYL2* in K9-derived mCherry-negative CMs treated with DMSO (NC) or 2C for 60 hours. Data are shown as mean ± SD (n=2 independent experiments with 4 replicates each). Two-way ANOVA with Šidák’s multiple comparisons test. ***P < 0.001, ****P < 0.0001. **(H-I)** Flow cytometry analysis of the percentage of mCherry-positive cells induced from mCherry-negative CMs by treatment with or without 2C for 60 hours. Data are shown as mean ± SD (n=3 independent experiments, represented as dots). **(J-K)** Flow cytometric plots showing EGFP-labeled CMs by lineage-tracing of K9-derived mCherry-negative CMs (**J**), and bar graph showing the percentage of mCherry-negative CMs expressing EGFP (**K**). Data are shown as mean ± SD (n=3 independent experiments). **(L)** Images showing the expression of ISL1 (red) and EGFP (green) in the cells induced from EGFP-positive/mCherry-negative CMs by treatment with or without 2C for 60 hours. DAPI (4′,6-diamidino-2-phenylindole) staining labeled nuclei as blue.

Additionally, we used a lineage-tracing system to unequivocally confirm the conversion of CMs to ISL1-expressing RCCs by 2C treatment. In this assay, CMs were transfected with plasmids encoding CreERT2 under the control of the TNNT2 promoter and an EGFP reporter following a flox-stop-flox cassette (Figure 3—figure supplement 3A). After optimizing viral titers and infection timing (Figure 3—figure supplement 3B-D), approximately 0.6% of K9-derived mCherry-negative CMs were permanently labeled with EGFP following 6 days of tamoxifen treatment (Figure 3J-K). These EGFP-labeled mCherry-negative CMs were then sorted by flow cytometry and treated with 2C for 60 hours. As expected, ISL1 expression was observed in these EGFP^+^ CMs (Figure 3L). Collectively, these results strongly indicate that the RCCs with ISL1 expression were genuinely generated by 2C treatment from TNNT2^+^ CMs.

## 2C-induced dedifferentiation of CMs provided a protective effect on cardiac infarction

To examine the reprogramming effect of 2C on CMs *in vivo*, we first tested 2C on primary neonatal rat CMs (Sakurai et al., 2014), which consistently induced RCC generation with corresponding ISL1 induction and morphological changes (Figure 4—figure supplement 1A). We then evaluated whether 2C could reprogram endogenous CMs to RCCs *in vivo*. Neonatal SD rats were intraperitoneally administrated 20 mg/kg of CHIR99021 and 10 mg/kg of A-485 daily for 5 days (Figure 4—figure supplement 1B). Compared to the vehicle/DMSO controls (NC), ISL1 was robustly induced in the CMs of rats treated with 2C without apparent effects on heart to body weight ratio (Figure 4—figure supplement 1C-D). These induced ISL1-expressing cells were widely distributed within the region 600 µm to 3000 µm down from the base of the hearts, including the aorta (Ao), left/right atria, and both ventricles (Figure 4—figure supplement 1E). All these ISL1^+^ cells also expressed TNNT2, indicating that 2C-induced RCCs originated from CMs *in vivo*. Importantly, neither CHIR99021 nor A-485 alone induced ISL1^+^ cells in ventricles, highlighting the combined effect of 2C on RCC induction (Figure 4—figure supplement 2).

**Figure 4.**
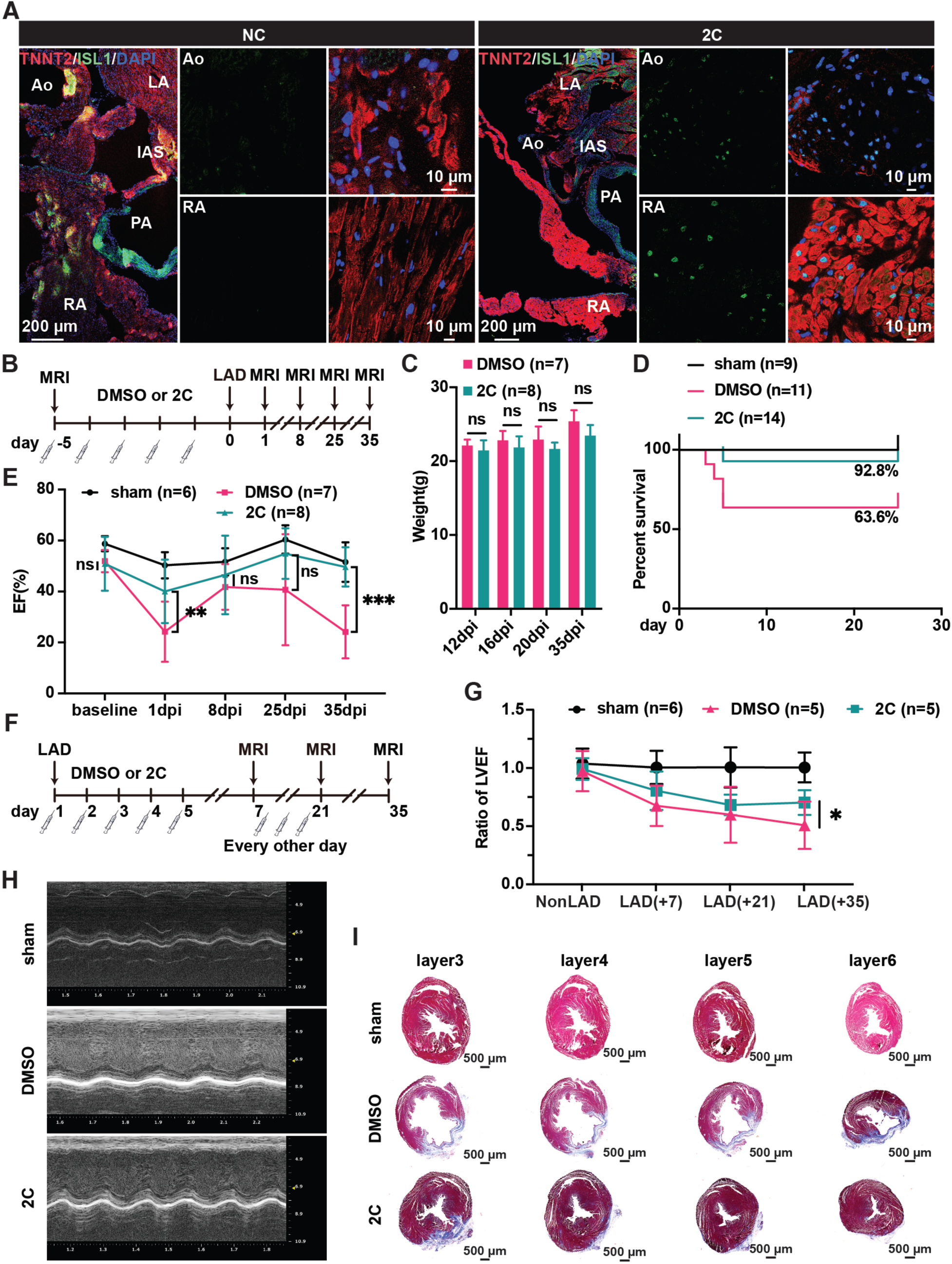
Heart regeneration *via* 2C-induced dedifferentiation of CMs. **(A)** Immunofluorescence staining of ISL1 (green) and TNNT2 (red) in cross-sectioned hearts from 2C or vehicle (DMSO)-treated (NC) adult 129SvJ mice. Ao, aorta. PA, pulmonary artery. LA, left atrial. RA, right atrial. IAS, interatrial septum. DAPI (4′,6-diamidino-2-phenylindole) staining labeled nuclei as blue. **(B)** Schematic illustration of the method used to examine the prophylactic effect of 2C in 129SvJ mice post MI. **(C)** Body weight of mice pre-treated with vehicle (DMSO) or 2C as shown in (**B**) at Day 12, Day 16, Day 20, and Day 35 after MI (dpi). Error bars represent SD. ns, not significant (P > 0.05). **(D)** Survival curve of sham-operated mice and mice pre-treated with vehicle (DMSO) or 2C as shown in (**B**), at indicated time points before or after MI. **(E)** Ejection fraction (EF) of sham-operated mice and mice pre-treated with vehicle (DMSO) or 2C as shown in (**B**), before MI (baseline) or at Day 1, Day 8, Day 25, and Day 35 after MI (dpi). Data are shown as mean ± SD. Two-way ANOVA with Tukey’s multiple comparisons test. ns, not significant (P > 0.05), **P < 0.01, ***P < 0.001. **(F)** Schematic illustration of the method used to examine therapeutic effect of 2C in the 129SvJ mice post MI. **(G)** Serial fMRI measurements showing the cardiac function from sham-operated mice and mice treated with vehicle (DMSO) or 2C at as shown in (**F**). Data are shown as mean ± SD. Two-way ANOVA with Tukey’s multiple comparisons test. *P < 0.05. **(H)** Echocardiography of sham-operated mice and mice treated with vehicle (DMSO) or 2C as shown in (**F**) at Day 35 post MI. **(I)** Masson staining of serial transverse sections of hearts from sham-operated mice and mice treated with vehicle (DMSO) or 2C as shown in (**F**) at Day 35 post MI.

Similarly, RCCs were efficiently induced in the Ao root and RA regions of the heart dissected from adult 129SvJ mice administrated 2C (20 mg/kg of CHIR99021 and 10 mg/kg of A-485) for 5 consecutive days (Figure 4A). To explore whether 2C-induced RCCs could rescue cardiac function in mice subjected to myocardial infarction (MI), we conducted both prophylactic and therapeutic studies. In the prophylactic setting, mice were pre-treated with either 2C or vehicle/DMSO for 5 days (Figure 4B) before inducing MI by ligation of left anterior descending (LAD) artery. When measuring cardiac function with magnetic resonance imaging (MRI) on days 1, 8, 25 and 35 post-MI, we found pretreatment with 2C significantly improved both cardiac function and survival rate without affecting body weight (Figure 4C-E). In the therapeutic setting (Figure 4F), mice treated with 2C post-MI displayed recovered cardiac function as assessed by left ventricular ejection fraction (LVEF) and echocardiography (Figure 4G-H). Cardiac fibrosis was largely ameliorated in 2C-treated mice, displaying significantly smaller scar sizes compared to control mice (Figure 4I). Altogether, these *in vivo* studies collectively indicate that 2C-induced RCCs possess regenerative capacities comparable to those generated *in vitro,* shedding light on the development of small molecule drugs with similar mechanisms for therapeutic use in repairing or regenerating hearts after cardiac injury.

## Dedifferentiation requirement of CHIR99021 and A-485

To elucidate the mechanisms behind the in 2C-induced dedifferentiation of CMs into RCCs, we performed bulk RNA-seq on K9-derived ISL1/mCherry-negative CMs treated with either DMSO as a negative control (NC) or the 2C combination for 60 hours (Figure 5A). Analysis of differentially expressed genes (DEGs) revealed substantial transcriptomic alterations; specifically, 2C treatment upregulated embryonic cardiogenesis genes such as MSX1, BMP4, TCF4, and LEF1, and downregulated genes associated with CMs (Figure 5B-C). Gene ontology (GO) analysis further indicated a suppression of genes involved in cardiac maturation and muscle contraction, contrasting with an enrichment in genes part of the catenin complex (Figure 5D-F). These findings were corroborated by qPCR, confirming the significant up-regulation of embryonic genes (e.g., MSX2, NKD1, PDGFC, CTNNA2) in K9-derived mCherry-negative CMs (Figure 5G).

**Figure 5.**
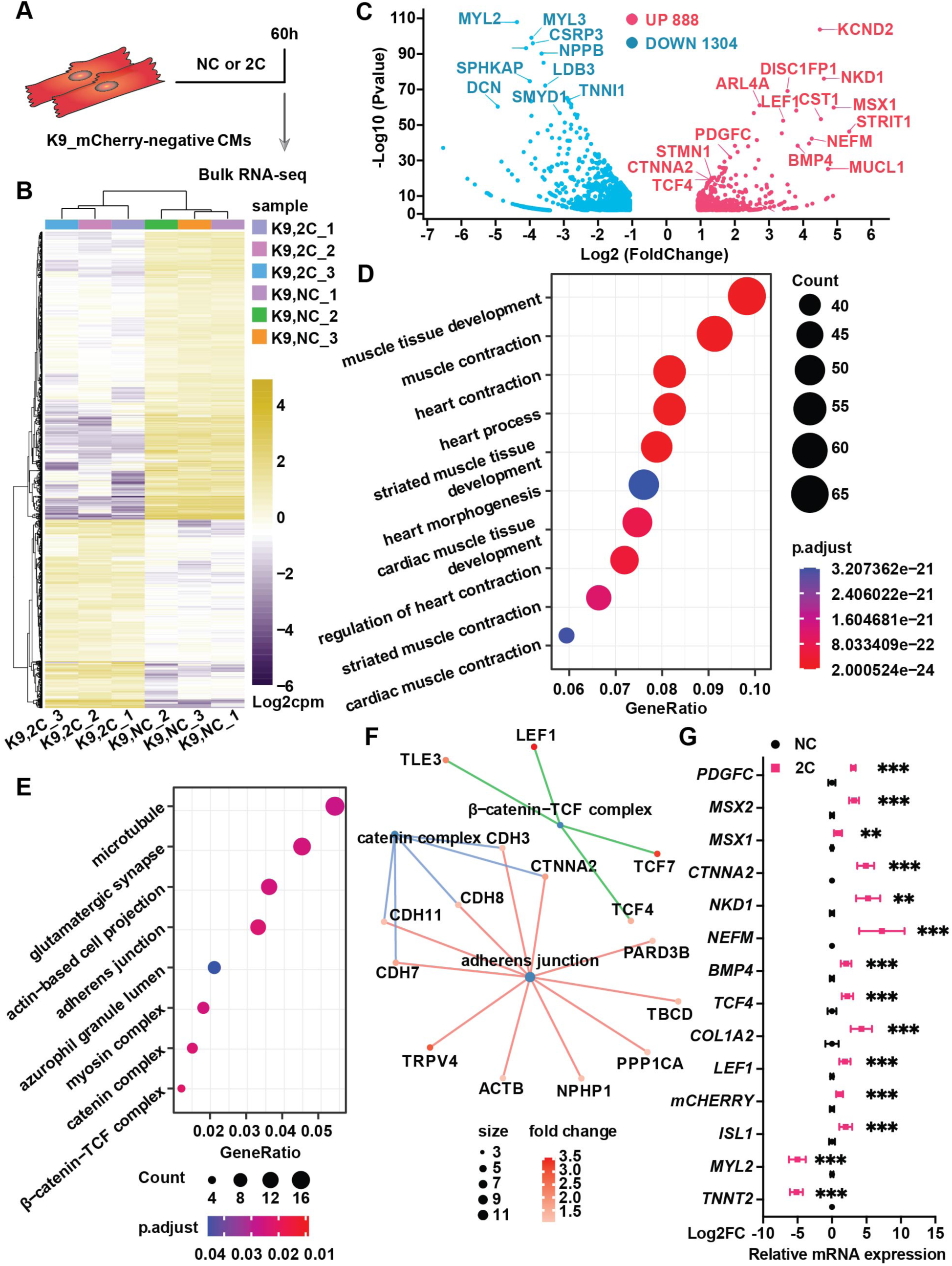
Bulk RNA-seq of analysis of 2C-treated ISL1/mCherry-negative CMs. **(A)** Scheme of bulk RNA-seq analysis of K9-derived mCherry-negative CMs with DMSO (NC) or 2C treatment for 60 hours. **(B)** Heatmap of differentially expressed genes (DEGs) in ISL1/mCherry-negative CMs treated with DMSO (NC) or 2C for 60 hours. **(C)** Volcano plot showing genes significantly changed by 60 hours of 2C treatment. **(D-E)** Gene Ontology (GO) analysis of downregulated (**D**) and upregulated (**E**) genes in ISL1/mCherry-negative CMs by 2C treatment for 60 hours, compared to DMSO (NC) treated cells. **(F)** Plotting GO terms of upregulated genes by 2C treatment with cnetlpot. **(G)** Relative expression fold-changes of indicated genes in K9-derived ISL1/mCherry-negative CMs by 60 hours of DMSO (NC) or 2C treatment. Data are shown as mean ± SD. Multiple unpaired t tests. **P < 0.01, ***P < 0.001.

We investigated the individual and combined effects of CHIR99021 and A-485 on inducing RCC characteristics. Morphological assessments showed that CHIR99021 significantly reduced both cytoplasmic and nuclear areas—a feature characteristic of RCCs—more effectively than A-485, which primarily reduced cytoplasmic area only (Figure 5—figure supplement 1A-C). Sarcomere disassembly was more pronounced with CHIR99021 treatment, aligning with its role in enhancing embryonic cardiogenesis genes like BMP4 and LEF1 (Figure 5—figure supplement 1D-E). A-485, while not affecting the nuclear area to the same extent, played a crucial role in the epigenetic reprogramming of CMs. It facilitated the suppression of CM-specific genes, such as TNNT2, through epigenetic modifications, enhancing the expression of RCC-specific genes such as ISL1 in conjunction with CHIR99021 (Figure 5—figure supplement 1D). This indicates that while CHIR99021 drives the initial dedifferentiation process by altering the cellular morphology and gene expression associated with embryonic cardiogenesis, A-485 enhances these effects by providing necessary epigenetic conditions for sustaining and stabilizing the RCC phenotype. Despite both compounds independently inducing reduced expression of TNNT2 and MYL2, as well as increased ISL1 expression, CHIR99021 uniquely influences a cohort involved in embryonic cardiogenesis, including BMP4, NKD1, MSX2, PDGFC, LEF1, and TCF4 (Figure 5—figure supplement 1D). Upon withdrawal of CHIR99021, A-485 or 2C, the cell percentage of ISL1^+^ cell was similarly and significantly reduced (Figure 5—figure supplement 1F). Notably, a drastic increase in the number of TNNT2^+^ CMs was observed only during the redifferentiation of 2C-treated cells, as confirmed by statistical analysis (Figure 5—figure supplement 1G). These findings suggest that while CHIR99021 plays a leading role in 2C-induced de-differentiation of CMs to RCCs, A-485 is also indispensable, particularly in modifying the epigenetic landscape to support the transition and maintenance of reprogrammed state.

## Reprogramming of CMs to RCCs by 2C went through an intermediate cell state

In the exploration of CM reprogramming to RCC using 2C treatment, single-cell RNA sequencing (scRNA-seq) was utilized to trace the progression through distinct cellular states of K9-derived mCherry-negative CMs treated with either DMSO (NC) or the 2C for 60 hours. Through UMAP analysis, we identified seven distinct clusters, observing notable increases in the proportions of cells in clusters 0, 2 and 3 following 2C treatment (Figure 6A-B). Cells within cluster 2 prominently expressed genes characteristic of RCCs, such as ISL1, BMP4 and FGF20—a gene crucial for the expansion of early embryonic progenitor cells (Cohen et al., 2007)), along with the cell proliferation marker MKI67 (Figure 6C-D). This distinct expression pattern marked them as transitioning towards an RCC phenotype. In contrast, clusters 0, 3, 4, and 5 maintained high expression levels of CM-specific markers like MYH6 and MYL2 (Figure 6C-D), indicating their retention of a mature CM identity. Interestingly, cells in clusters 1 and 6 exhibited characteristics of intermediate cells (ICs), expressing both dedifferentiation markers such as ACTA2 and genes linked to CM development, including COL1A1, COL1A2, and COL3A1 (Cui et al., 2020; Mononen et al., 2020) (Figure 6C-D). This highlighted these clusters as transitional states in the reprogramming process from CMs to RCCs.

**Figure 6.**
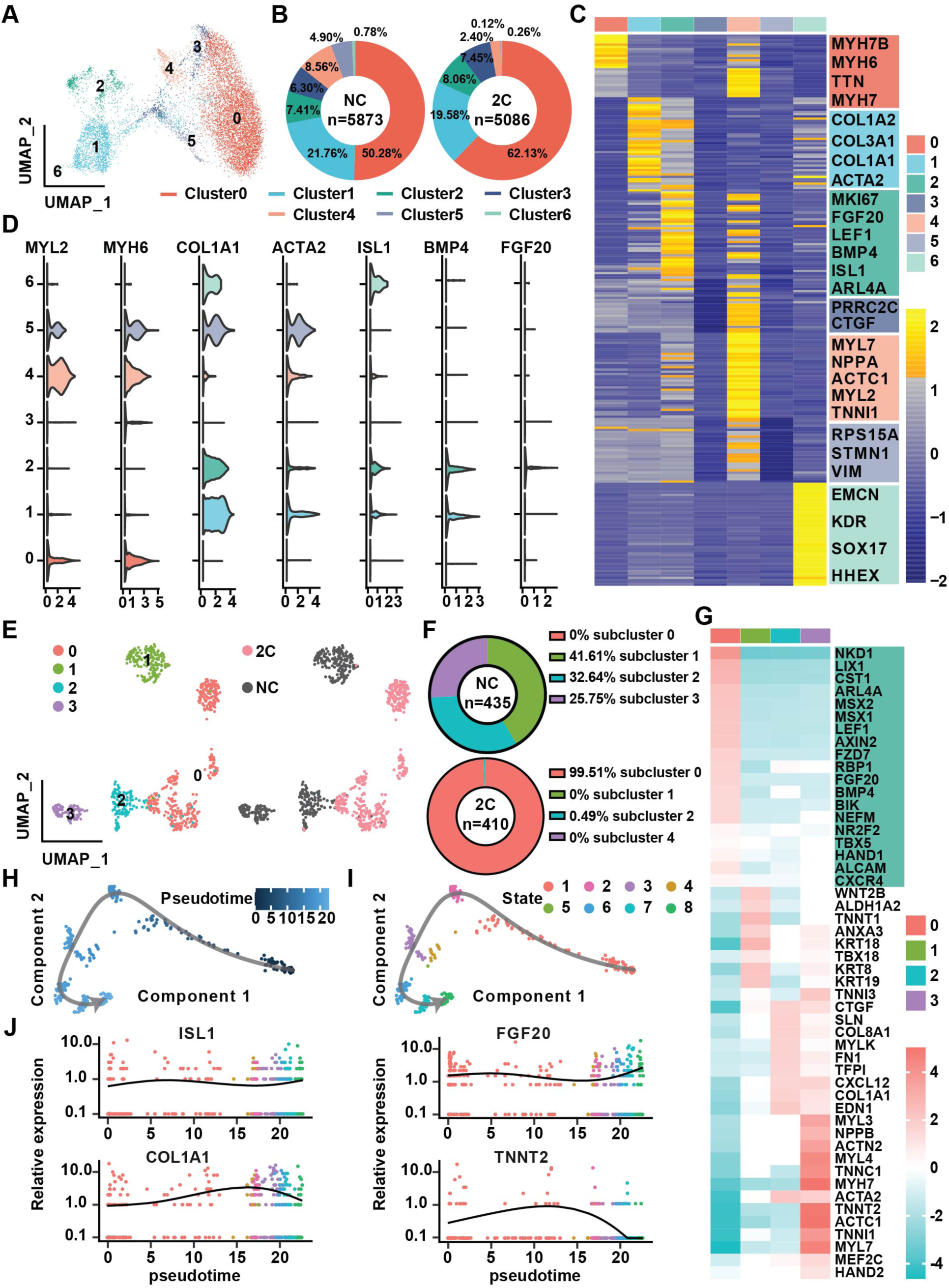
Single-cell RNA-seq of 2C-treated mCherry-negative CMs. **(A)** UMAP analysis showing 7 clusters in cells induced from K9-derived mCherry-negative CMs by treatment with DMSO (NC) and 2C for 60 hours. **(B)** The percentage of cells in the 7 indicated clusters, following DMSO (NC) or 2C treatment. **(C)** Heatmap showing the differentially expressed genes in the cells from 7 indicated clusters. The representative marker genes of 7 indicated clusters were listed on the right. **(D)** Violin plots showing the expression levels of marker genes of CMs (MYL2, MYH6), ICs (COL1A1, ACTA2), and RCCs (ISL1, BMP4, FGF20) among cells from 7 indicated clusters. **(E)** UMAP analysis showing the second-level clustering of cluster 2 into 4 subclusters (left), which exhibited dramatic distinction under condition of 2C or NC (right). **(F)** The percentage of cells in the 4 indicated subclusters within cluster 2, following DMSO (NC) or 2C treatment. **(G)** Heatmap showing the differentially expressed genes among cells from 4 subclusters of cluster 2. Genes related to RCCs are highlighted in the green blocks on the right. **(H-I)** Pseudotime trajectory showing changes across various cell states upon 2C treatment, which were presented with different developmental pseudotime points (**H**) and cell states (**I**). **(J)** Curves showing the dynamic expression of representative genes of RCCs (ISL1, FGF20), ICs (COL1A1), and CMs (TNNT2) along indicated pseudotime points.

Further detailed analysis of cluster 2, comparing 435 cells from DMSO-treated samples with 410 cells from 2C-treated samples, allowed us to identify a prominent subcluster (subcluster 0) comprising 408 cells (48.3% of the cluster), exclusively found in 2C-treated samples (Figure 6E-F). Cells in this subcluster were defined by high expression of key embryonic cardiogenesis genes such as MSX1, LEF1, BMP4, MSX2, and HAND1, along with genes typically co-expressed with ISL1 in SHF progenitors, including NR2F2, TBX5, ALCAM (Ghazizadeh et al., 2018), and CXCR4 (Andersen et al., 2018) (Figure 6G). This specific gene expression profile robustly indicates that the cells in subcluster 0 have adopted an RCC identity, indicative of successful induction by 2C treatment. Thus, a unique gene set that included LIX1, NKD1, PDGFC, ARL4A, AXIN2, FZD7, ISL1, MSX1, MSX2, BMP4, LEF1, and HAND1 was found to determine RCC state during the reprogramming of CMs by 2C, and hence designated as RCC genes.

To further quantify this transition, pseudo-time analysis was performed, clearly delineating the 2C-induced reprogramming trajectory from CMs, highly expressing TNNT2, through ICs, characterized by the expression of COL1A1, to RCCs, marked by elevated levels of ISL1 and FGF20) (Figure 6H-J). This analysis confirms the dynamic and stepwise nature of cellular transformation under 2C treatment.

## Epigenetic regulation of CMs and RCCs genes by 2C treatment

As previously documented (Lasko et al., 2017), A-485 functions as a p300 acetyltransferase inhibitor, specifically reducing acetylation at lysine 27 of histone H3 (H3K27Ac), but not at lysine 9 (H3K9Ac), as shown in studies with PC3 cells. Our data corroborate these findings, showing a notable decrease in H3K27Ac levels in CMs treated with A-485 alone, while levels of H3K9Ac remain unaffected (Figure 7— figure supplement 1A-E). However, treatment with CHIR99021 or the combined 2C regime did not significantly alter the acetylation patterns of these histone marks (Figure 7—figure supplement 1). Intriguingly, we observed that although the number of H3K9Ac^+^ cells remained unchanged in CMs, the fluorescence intensity was significantly decreased after A-485 treatment (Figure 7—figure supplement 1F), suggesting a distinctive regulatory role of A-485 in modulating histone acetylation in CM reprogramming.

**Figure 7.**
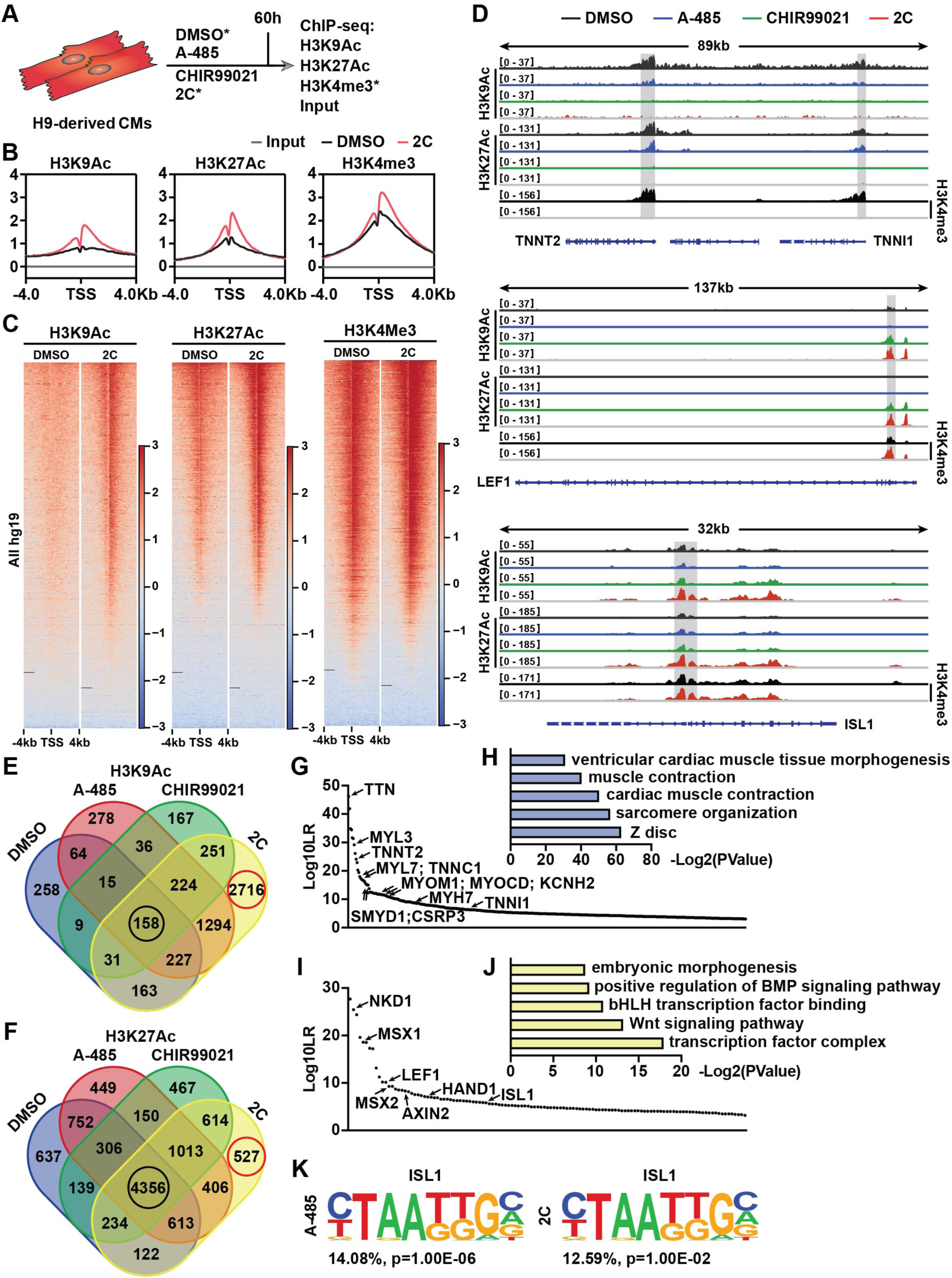
Chromatin immunoprecipitation-sequencing (ChIP-seq) analyses of chemical-treated H9 hESC-derived CMs. **(A)** Schematic illustration of ChIP-seq analysis of H9-derived CMs subjected to DMSO, A-485, CHIR99021, or 2C treatment for 60 hours. **(B)** Average ChIP-seq signal profiles showing the indicated histone modifications around the TSS in the input and ChIP samples prepared from DMSO and 2C-treated cells. **(C)** Heatmap showing the whole-genome wide distribution of H3K9Ac, H3K27Ac, and H3K4me3 peaks within a range of ±4kb from TSSs in the cells treated with DMSO or 2C for 60 hours. **(D)** H3K9Ac and H3K27Ac peaks surrounding CM genes (TNNT2 and TNNI1) and RCC genes (LEF1 and ISL1) in the cells treated with DMSO, A-485, CHIR99021 or 2C for 60 hours, and H3K4me3 peaks surrounding the same genes in the cells treated by DMSO or 2C. The y-axis represents the number of counts. **(E-F)** Veen diagram showing the number of annotated genes with H3K9Ac or H3K27Ac enrichment in the cells treated with DMSO, A-485, CHIR99021 or 2C for 60 hours. Red circles indicate the number of genes with unique H3K9Ac or H3K27Ac enrichment induced by 2C treatment; black circles indicate the number of genes with H3K9Ac or H3K27Ac enrichment unaffected by any chemical treatment. **(G-J)** The annotated genes with the most significant changes in H3K9Ac enrichment following treatment with DMSO (**G**) or 2C (**I**) were ranked by Log10LR and analyzed by GOs (**H** and **J**), respectively. **(K)** ISL1 binding motifs identified from the cells treated with A-485 or 2C.

To explore the molecular mechanisms by which the 2C treatment modulates gene expression during the reprogramming of hESC-derived CMs into RCCs, we conducted chromatin immunoprecipitation followed by sequencing (ChIP-seq) analyses. This study focused on H3K27Ac and H3K9Ac modifications across various treatment conditions (Figure 7A). Our analysis identified regions near transcription start sites (±4 kb) that were rich in H3K4me3, indicative of active promoters in both control (DMSO-treated) and 2C-treated cells (Figure 7B-C). Following 2C treatment, there was a marked increase in H3K9Ac and H3K27Ac levels around the transcription start sites (TSS), suggesting enhanced gene activity. Specifically, 2C treatment significantly increased H3K9Ac enrichment at 4,485 gene promoters and H3K27Ac at 2,560 gene promoters, with corresponding decreases in 346 and 1,834 genes, respectively (Figure 7—figure supplement 2A-F). Notably, genes that showed reduced acetylation in response to A-485 also displayed similar trends under 2C treatment, highlighting genes with functions crucial to cardiac muscle activity such as contraction and sarcomere organization (Figure 7—figure supplement 2). In comparison with treatments using A-485 or CHIR99021 alone, the 2C combination notably augmented acetylation at genes involved in cell cycle regulation and cell division, suggesting a synergistic effect of the dual treatment in promoting gene activation essential for cardiac regeneration (Figure 7—figure supplement 2).

We also specifically analyzed the acetylation peaks at promoters of key CM genes and RCC genes. A-485 treatment predominantly decreased H3K9Ac and H3K27Ac peaks at CM gene promoters such as TNNT2, TNNI1, MYL7, MYH6, and MYH7, aiding their downregulation. These reductions were intensified under 2C treatment, reflecting its potent effect in repressing mature cardiomyocyte markers while activating regenerative pathways (Figure 7D and Figure 7—figure supplement 3). In contrast, CHIR99021 effectively upregulated H3K9Ac and H3K27Ac at promoters of the RCC genes such as LEF1, AXIN2, BMP4, LIX1, MSX1, MSX2, and NKD1, with 2C treatment further enhancing these effects, confirming its robust impact on inducing a regenerative phenotype in treated cells (Figure 7D and Figure 7—figure supplement 3).

Moreover, the distribution of H3K9Ac and H3K27Ac changes during 2C-induced dedifferentiation revealed that these modifications are more dynamically associated with H3K9Ac, particularly in the transition of CMs to RCCS. This dynamic regulation was illustrated by the significant number of genes showing exclusive enrichment under 2C treatment compared to control or single-drug treatments (Figure 7E-F). In detail, of genes with H3K9Ac enrichment, 2716 (or 46.1%) out of 5891 annotated genes were only enriched in the cells treated with 2C, while only 527 (or 4.9%) out of 10785 annotated genes with H3K27Ac enrichment were exclusively observed (Figure 7E-F). To further confirm this notion, we performed a comparative analysis of annotated genes between the cells treated with 2C or DMSO to validate the most significant differential genes (i.e., Log10LR>3) under the respective condition. In comparison to 430 genes with H3K9Ac enrichment observed in DMSO-treated cells, which were mainly involved in cardiac muscle contraction (Figure 7G-H), 2C treatment markedly enhanced the enrichment of H3K9Ac in 123 genes associated with the transcription factor complex and Wnt signaling pathway (Figure 7I-J). In particular, 12 CM genes, including TNNT2, were ranked in the top 90 out of 430 annotated genes with H3K9Ac enrichment. In contrast, 7 RCC genes, including ISL1, were ranked in the top 50 out of 123 annotated genes with H3K9Ac enrichment.

Lastly, our findings were substantiated by DNA-binding motif analyses, which showed increased accessibility of ISL1 binding sites only in cells treated with A-485 or the 2C combination, underscoring the critical role of A-485 in activating RCCs genes through ISL1-dependent mechanisms (Figure 7K). Collectively, these results underscore the efficiency of combining CHIR99021 and A-485 in reprogramming CMs to RCCs, opening new avenues for developing therapies aimed at cardiac regeneration.

## Discussion

The generation of regenerative cardiac cells (RCCs) from cardiomyocytes (CMs) using the 2C treatment, a combination of CHIR99021 and A-485, represents a significant advancement in cardiac regenerative medicine. This study demonstrates that the synergistic interaction between these two molecules is essential for the reprogramming of endogenous CMs into RCCs, as evidenced by the absence of RCCs in postnatal rat hearts treated with either molecule alone (Figure 4—figure supplement 2). CHIR99021, a well-documented GSK3 inhibitor, activates the Wnt signaling pathway, thereby promoting the proliferation of human pluripotent stem cell-derived CMs (Buikema et al., 2020; Quaife-Ryan et al., 2020). It also induces H3K27Ac at promoters of critical cardiac genes such as TNNT2, MYH6, and ISL1 (Quaife-Ryan et al., 2020). This observation aligns with our findings, which show a slight increase H3K27Ac levels in CMs following CHIR99021 treatment (Figure 7D and Figure 7—figure supplement 1). Moreover, our results reveal that the p300/CBP inhibitor A-485 facilitates CM dedifferentiation into RCCs by epigenetically suppressing CM-specific gene expression (Figure 7D), consistent with a decline in the fluorescence intensity of H3K27Ac and H3K9Ac in CMs (Figure 7—figure supplement 1C and E). Concurrently A-485 enhances modifications conducive to the RCC state, such as H3K9Ac and H3K27Ac on genes like ISL1 (Figure 7D). This dual mechanism of action underscores the potential of utilizing epigenetic modifiers like A-485 to influence cell fate decisions in terminally differentiated cells, suggesting broader applications in regenerative biology.

The RCCs induced by the 2C treatment not only re-expressed genes essential for embryonic cardiogenesis but also demonstrated the capability to differentiate into three cardiac lineages *in vitro*, mirroring the properties of ISL1^+^ second heart field cardiac progenitors (Bu et al., 2009; Moretti et al., 2006). Although these 2C-induced RCCs show limited proliferation compared to natural ISL1^+^ progenitors, their potential for expansion under defined conditions presents an intriguing avenue for future research. These insights contribute significantly to the understanding of cardiac lineage plasticity and highlight the therapeutic potential of induced multipotent cardiovascular progenitors for heart repair and regeneration.

Despite these promising findings, several limitations warrant further investigation. The modest proliferation ability of 2C-induced RCCs relative to natural cardiac progenitors suggests that optimizing the culture conditions or treatment protocols may enhance their regenerative capacity. Additionally, the specific molecular mechanisms through which CHIR99021 and A-485 synergistically promote RCC formation remain incompletely understood. Future studies should aim to delineate these pathways more clearly, potentially through *in vivo* lineage tracing to map the fate decisions of individual cells. Moreover, the potential off-target effects of 2C treatment and its long-term impacts on cardiac function and structure need comprehensive evaluation. As we advance, exploring the scalability of this approach and its applicability to other types of terminally differentiated cells could broaden the scope of regenerative therapies, offering new strategies for numerous degenerative conditions. In conclusion, this study not only sheds light on the cellular and molecular intricacies of cardiac cell dedifferentiation but also opens new pathways for the development of regenerative therapies aimed at heart disease.

## Acknowledgments

This work is supported by the National Natural Science Foundation of China (32030031 to S.D.), Beijing Natural Science Foundation (JQ22016 to T.M.), the National Key R&D Program of China (2022YFA1103704 to S.D.; 2022YFA1104503 to Y. N.), Center for Life Sciences (to S.D.).

We also thank the Center for Pharmaceutical Technology, Tsinghua University for the activity screening platform, Biomarker Technologies Corporation, Beijing, China and BeiJing Geek Gene Technology Co Ltd for technical support, and support from Tsinghua-Peking Center for Life Sciences.

## Author Contributions

Conceptualization, S.D. and W.Z.; Methodology, W.Z., H.L., T.H.M. and Y.N.; Investigation, W.Z., C.Y.W., K.Z.H., P.Q.W., B.W.W., and H.G.; Visualization, W.Z., P.Q.W., B.W.W., H.G., and D.W.; Funding Acquisition, S.D., T.H.M., and Y.N.; Project Administration, W.Z., H.L., T.H.M., and Y.N.; Supervision: S.D., and W.Z.; Writing – Original Draft, W.Z., T.H.M., and S.D.; Writing – Review & Editing: W.Z., T.H.M., Y.N., S.D.

## Data and Materials availability

All targeted amplicon sequencing data have been deposited in the Sequence Read Archive of the NCBI under the BioProject accession number <u>PRJNA903530.</u> All data are available in the main text or the supplementary materials.

