## Supplemental File for "Pharmacologically inducing regenerative cardiac cells by small molecule drugs"

**This PDF file includes:**

Supporting text
Figure1—figure supplement 1-2
Figure2—figure supplement 1
Figure3—figure supplement 1-3
Figure4—figure supplement 1-2
Figure5—figure supplement 1
Figure7—figure supplement 1-3

Table 1
SI References

**Other supporting materials for this manuscript include the following:**

Movies 1-2

### Supporting Information Text

#### Materials

**Mice.** All animal experiments in this study were performed in accordance with the guidelines and regulations of IACUC (Institutional Animal Care and Use Committee) of Tsinghua University, Beijing, China. All the animal protocols used in this study have been approved by IACUC of Tsinghua University. All mice were housed in individually ventilated cages (maximal six mice per cage) at Tsinghua University. The mice/rats were maintained on a 12/12-h light and dark cycle, and at 22–26°C with sterile pellet food and water ad libitum. The adult 129SvJ mice and pregnant SD1 rats at gestational day 17.5 were purchased from Beijing Vital River Laboratory Animal Technology. All adult mice were 8-12 weeks old when used for experiments.

### Methods

**Drug administration.** CHIR99021 (20 mg/kg, selleck) and A-485 (10 mg/kg, MCE) were administration in SD1 neonatal ( at day of birth, P1) rats and 129SvJ adult mice by intraperitoneal (i.p.) injection as indicated. Administration with equal volume of DMSO was used as negative control.

**Chemically defined induction of human embryonic stem cells differentiation into cardiomyocytes.** H9 and HUES7 human embryonic stem cells (hESCs) (WiCell) were maintained in mTeSR1 medium (Stem Cell Technologies) on Matrigel (BD Biosciences) coated plates, according to the manufacturer's instructions. An established protocol (Lian et al., 2013; Tohyama et al., 2013) was modified for cardiac differentiation of hESCs. In brief, hESCs were seeded into the plates at $1.5 \times 10^5$ cells/cm$^2$. When reaching 90%-95% confluency, cells were cultured with RPMI/B27$^{-INS}$ media (the RPMI 1640 basal medium (Gibco) with 1 × B27 minus insulin supplement (Gibco)) containing 20 ng/ml hActivin A (R&D), 20 ng/ml hBMP4 (R&D), and 1.5 μM CHIR99021 (Selleck) for 2 days to initiate differentiation. Subsequently, the culture media were changed to RPMI/B27$^{-INS}$ media containing 5 μM IWP2 (Tocris) for 3 days and then changed to RPMI/B27$^{-INS}$ media for another 3 days. From day 8, the cardiomyocyte (CM) culture media RPMI/B27 was used to culture cells for 6 days, with changes every 2-3 days during this period. Contracting CMs were observed as early as day 10 of differentiation. To purify the CMs, cells at day 14 of differentiation were cultured in CM selection media (glucose-free DMEM (Gibco) supplemented with 4 mM lactate (Sigma)) for 4-6 days. CM selection media was changed every 2 or 3 days for eliminating dead cells. Cells were then digested by TrypLE (Gibco), reseeded onto fibronectin (BD Biosciences) coated plates with media containing MEM-a (Gibco), 5% FBS (Gibco) and 5 μM Y-27632 (Selleck) for 24 hours and recovered in CM culture media for 4 days.

**Establish ISL1$^{mCherry/+}$ hESC lines.** Knock-in of ISL1-mCherry reporter was performed in H9 and HUES7 hESC lines using CRISPR-Cas9 system (Ran et al., 2013). In details, three single-guide RNAs were designed and cloned into the PX459 plasmid (Addgene). The reporter gene mCherry was placed directly downstream of ISL1 start condon, and followed by the hygromycin resistance gene flanked by Loxp sites. All DNA fragments were contructed into the pEGFP-N3 vector (Addgene) from BamHI and NotI sites using NEBuilder® HiFi DNA Assembly Master Mix (New England BioLabs) according to the manufacturer's instructions. Both CRISPR-Cas9 plasmids and trageting vectors were delivered into hESCs using P3 Primary Cell 4D-Nucleofector™ X Kit (Lonza) according to the manufacturer's instructions. Subsequenctly, cells with knock-in of ISL1-mCherry reporter were selected by hygromycin (500 ng/ml) (Thermo Fisher Scientific) and identified using PCR (Primer F: GGGCCCGGGATCCGAAGGAAGAGGAAGA, Primer R: GAGGTCGA GATCCTAAGCTTGGC). To excise the selection cassette, transient expression of Cre recombinase was performed by transfection using Lipofectamine® 3000 (Life Technologies), followed by flow cytometry-based sorting of EGFP-positive cells. The established ISL1$^{mCherry/+}$ hESC clones were genotyped by PCR (Primer F: CCGCGGGCCCGGGATCCGTCA GTCCGCGGAGTCAAC, Primer R: TAGAGTCGCGGCCGCGCCGCAACCAACACATA) and sequencing.

**Chemical induction of regenerative cardiac cells from cardiomyocytes.** CMs were seeded onto culture well, fibronectin-coated 6-well-plates at a density of 1.6 × $10^5$-3.0 × $10^5$ cells/$cm^2$ and cultured in CM culture media. When the CMs recovered contraction, the culture medium was changed with chemically reprogramming medium (MCDB131 (Gibco), 25 mg/L $NaHCO_3$ (Sigma), 4 mM Glucose (Sigma), 1% Clutmine (Gibco), 1 × N-2 supplement (Gibco), and 20% 1 × B-27 supplement (Gibco), 10 μM CHIR99021 (Selleck) and 0.5 μM A-485 (MCE). Cells were induced to reprogram for 24 hours, 48 hours or 60 hours, depending on experiments. Cells treated with equal volume of DMSO was used as negative control.

**Neonatal cardiomyocytes isolation.** Neonatal cardiomyocytes (at postnatal day 3 or P3) were isolated following a previously reported protocol (Sakurai et al., 2014) with a slight modification. Briefly, the hearts were collected into 50 ml BD tubes containing 30 ml ice-chilled CMF-HBSS (Gibco) on ice. The rinsed hearts were transferred to a 10-cm plastic dish (CORNING) placed on ice and scissored after removal of large vessels and/or unwanted tissues. After adding frozen trypsin (Gibco) at a final concentration of 50 μg/ml, the dishes were sealed with parafilm and placed overnight at 4°C. On the next day, 5% FBS (Gibco) was added and incubated at 37°C for 30 minutes. After the addition of Leibovitz L-15 (Gibco) containing collagenase (Gibco), dishes were gently shaken for 45 minutes. When the tissue was homogenized, the cells were pipetted, filtered through 70 μm cell strainer (NEST), and transferred into a new 50 ml BD tube. Residual cells were resuspended with 5 ml of fresh Leibovitz L-15, filtered through 70 μm cell strainer, and transferred as well. Cells were further digested at 37°C for 40 minutes and then centrifuged at 200 × g for 5 minutes. After re-suspension with 10 ml DMEM (Gibco) containing 10% FBS and 20 U/ml penicillin/streptomycin (Gibco), cells were transferred into 10-cm plastic dish and incubated at 37°C for 1 hour. Supernatant containing the cardiomyocytes was collected and centrifuged at 200 × g for 5 minutes. Cardiomyocytes were resuspended with DMEM containing 10% FBS and 10 U/ml penicillin/streptomycin, counted and seeded on laminin-coated 24-well plates at a density of 5 × $10^5$ cells/$cm^2$. On the next day, the media was changed to DMEM/MEM (Gibco) containing 5% FBS, 10 U/mL penicillin/streptomycin and 0.1 mM BrdU (Sigma). After 48 hours, the media was changed to DMEM/MEM containing 5% FBS and 10 U/ml penicillin/streptomycin and the fresh media was changed every 2-3 until initiation of chemical treatment.

**Myocardial infarction model.** Myocardial infarction (MI) was performed in adult mice at age of 9-12 weeks by left anterior descending coronary artery (LAD) ligation as previously described (Mahmoud et al., 2014). In details, adult mice were anesthetized by intraperitoneal injection of tribromoethanol (Sigma-Aldrich, 200 mg/kg) and artificially ventilated with tracheal intubation. Lateral thoracotomy at the 3rd-4th intercostal space was performed by blunt dissection of the intercostal muscles following skin incision. The LAD was ligated by a 7/0 non-absorbable silk suture. The successful ligation of the LAD was verified by visual inspection of the apex color turning to bloodless. Following ligation, the thoracic wall incisions and the skin wounds were sutured with a 4/0 non-absorbable silk suture. Mice were warmed for several minutes until recovery. In the sham controls, we performed the same procedures without LAD ligation.

**Small molecule libraries.** To identify compounds that enable induction of ISL1-expressing cells from CMs, we performed a large-scale screening of a collection of 235 small molecules modulating signaling pathways, epigenetic modifications, metabolites and nuclear receptors involved in the cardiac development (Supplementary Table 1), and two commercial compound libraries, Sigma-LOPAC (Sigma) and Selleck-FDA (Selleckchem). 5 μM or 0.5‰ DMSO was used as negative control in compound screening. Small molecules in our collection were purchased from Sigma, Tocris Bioscience, Selleck, and MedChemExpress.

**Immunocytochemistry and immunohistochemistry.** For immunocytochemistry, cells were fixed with 4% paraformaldehyde (Sigma), blocked in PBS (Gibco) containing 4% FBS (Gibco) and 0.3% Triton X-100 (Sigma) at room temperature for 30 minutes and incubated with the primary antibody diluted in PBS containing 5% skim milk (AMRESCO) and 0.1% Triton X-100 at 4°C overnight. After washing with PBS, cells were incubated with secondary antibodies conjugated with Alexa Fluor®-488 or -549 (Life Technologies) diluted in PBS at room temperature for 1 hours and stained with 4,6-diamidino-2-phenylindole (DAPI, Sigma) for 5 minutes. For immunohistochemistry, harvested hearts were fixed with 4% paraformaldehyde at 4°C for 1 hour and further dehydrated in 30% sucrose at 4°C overnight. Next, fixed hearts were embedded in optimal cutting temperature compound (SAKURA) and sectioned at 10 μm thickness with a cryostat (Leica CM1900). Immunostaining of cardiac tissue sections were performed following the procedure of immunocytochemistry described above. Quantitative analysis of the immunostained samples were captured and analyzed by Opera Phenix (PerkinElmer). The following antibodies were used: OCT4 (Santa Cruz Biotechnology; 1:400), SOX2 (Abcam; 1:400), NANOG (Abcam; 1:400), SSEA1 (Stemgent; 1:200), MESP1 (ASB, 1:200), MYL2 (Abcam; 1:500), mCherry (Abcam; 1:200), GATA4 (Abcam; 1:500), ISL1 (Developmental Studies Hybridoma Bank; 1:100), NKX2-5 (Santa Cruz Biotechnology; 1:200), TBX5 (Sigma, 1:200), SMA (Sigma, 1:500), TNNT2 (Sigma, 1:1000), GFP (Sigma, 1:500), CD31 (BD Biosciences; 1:200), VEcadherin (R&D; 1:200), MEF2C (Cell Signaling Technology; 1:200), NR2F2 (Cell Signaling Technology; 1:200), Phospho-Histone H3 (Cell Signaling Technology; 1:200), SSEA4 (R&D; 1:200), Smooth Muscle Actin-a (Thermo Scientific; 1:500). Images were captured using a confocal Zeiss LSM710 and Olympus IX83 inverted microscope. Trichrome staining was performed using the Trichrome Stain (Masson) Kit (Sigma, HT15).

**Flow-cytometry analysis and fluorescence-activated cell sorting (FACS).** Cells subjected to flow-cytometry analysis were harvested and dissociated by using Versene (Gibco, used on D6) or TrypLE (Gibco, used on SD4). For direct flow-cytometry analysis of surface proteins, samples were incubated with Zombie VioletTM (eBioscience) diluted at 1:500 in PBS (Gibco) for 15 minutes in the dark. After washing once with PBS containing 2% FBS (Gibco), samples were stained with antibodies and analyzed using FACS. For analysis of intracellular proteins, cells were fixed and permeabilized with reagents in the Foxp3 Staining Buffer Set (eBioscience), blocked in PBS containing 2% FBS, and incubated with primary antibodies against ISL1 (Developmental Studies Hybridoma Bank; 1:200), TNNT2 (Sigma, 1:1000), mCherry (Abcam; 1:200). Isotype-matched normal IgGs (Life Technologies) served as negative controls.

After stained with secondary antibodies conjugated with Alexa Fluor®-488 or -549 (Life Technologies), cells were analyzed and quantified by flow cytometry using a BD FACSAria III Cell Sorter (BD Biosciences). To sort K7 or K9-derived mCherry$^+$ cells, mCherry-negative CMs or TNNT2-EGFP$^+$ CMs in this study, corresponding living cells were harvested and incubated with Zombie VioletTM (eBioscience) at 1:500 in PBS for 15 minutes in the dark. After washing with PBS containing 2% FBS, cells were resuspended with CM culture media and subject to FACS using the BD FACSAria III Cell Sorter.

**RNA extraction, reverse-transcription PCR and quantitative real-time PCR (qRT-PCR).** Total RNA extracted from $>10^5$ cells with AxyPrep Multisource RNA Miniprep Kit (AXYGEN Biosciences), according to manufacturer's instructions, or extracted from $<10^5$ cells using TRIzol (Gibco). After elimination of genomic DNA by DNaseI, cDNA was synthesized by reverse transcription of 1 μg total RNA with iScript cDNA Synthesis Kits (Bio-Rad). Quantitative RT-PCR was performed and analyzed with iQ SYBR Green Supermix (Bio-Rad) using CFX384 Touch Real-Time PCR Detection System (Bio-Rad). Relative mRNA expression of specific genes was normalized to the level of human *GAPDH* transcripts.

**Western blot.** Cells were lysed on ice in RIPA buffer (50 mM Tris-HCl pH 7.5, 150 mM sodium chloride, 0.25% sodium deoxycholate, 0.1% Nonidet P-40, and 0.1% Triton X-100) containing protease and phosphatase inhibitors (Roche). The homogenized samples were centrifuged at 14,000 rpm for 5 min at 4°C, and the supernatant of each sample was transferred into a new respective tube and quantified by Bradford Assay. After addition of 5 × SDS sample buffer (50 mM Tris-HCl PH 7.5, 150 mM NaCl, 0.25% Sodium deoxycholate, 0.1% Nonidet P-40, 0.1% Triton X-100, 5% β-mercaptoethanol), samples were boiled for 5 minutes. Next, 10 μg of total protein from each sample was subjected to SDS-polyacrylamide gel electrophoresis (SDS-PAGE) to separate proteins based on their different molecular weight. Western blot was performed using antibodies of β-Actin (Santa Cruz Biotechnology; 1:500), and ISL1 (Developmental Studies Hybridoma Bank; 1:200). The results were visualized by AI600 (GE), and quantitatively analyzed by ImageJ.

**Lentiviral transduction.** All lentiviral shuttle plasmids were constructed with NEBuilder® HiFi DNA Assembly Master Mix (New England BioLabs) according to the manufacturer's instructions. Briefly, DNA fragments with ~20-bp overlapping ends were generated by restriction enzyme digestion or PCR. Then DNA fragments were mixed with the Assembly Master Mix at a volume ratio of 1:1 and incubated at 50°C for ~30 minutes. The products were transformed into Trans1-T1 Phage Resistant Chemically Competent Cell (TransGen Biotech). The TNNT2 promoter was amplified from genomic DNA by PCR (Primer F: GTCATGGAGAAGACCCACCTT, Primer R: GATCCTGGAGGCGTCTGC). All of the plasmids used in this study were purchased from Addgene. After package of lentivirus in HEK293T cells, the concentrated lentiviral supernatants were mixed with fresh CM culture media at a volume ratio of 1:1 and used to transduce isolated CMs for 48 hours. Next, CMs were cultured with CM culture media containing 2 μM Tamoxifen (Selleck) for another 6 days. The EGFP$^+$ CMs sorted by FACS were used in 2C-induced reprogramming.

**Multibarcode RNAseq.** K9-derived mCherry-negative CMs sorted by FACS were treated with DMSO or 2C for 60 hours and then prepared for Smart-Seq2. The RNA-seq libraries were generated from these samples according to a previously reported Smart-Seq2 protocol with minor modifications (Picelli et al., 2013). Briefly, cells were lysed in lysis buffer containing RNase inhibitor (TaKaRa). RNAs were captured with 25 nt oligo (dT) primers and reversed into cDNAs. After amplification and purification, cDNAs were sheared to approximately 300 bp by CovarisS2 and captured by DynabeadsR MyOne Streptavidin C1 beads (Thermo Fisher). All libraries were constructed using a Kapa Hyper Prep Kit (Kapa Bio-systems) and sequenced on 150 bp paired-ends Illumina Novaseq 6000 platform.

**Single-cell RNA sequencing (scRNA-seq) analysis.** After sorted by FACS, K9-derived mCherry-negative cells were treated with DMSO or 2C for 60 hours, and then subject to scRNA-seq. The single-cell suspensions were prepared in PBS containing 0.04% BSA (Sigma). The scRNA-seq libraries were constructed by Chromium Next GEM Single Cell 3' Kit v3.1 up at Chromium Controller (10 × Genomics) and the quality of these libraries were assessed by Qubit 4.0 and the Agilent 2100. Sequencing was performed on the Illumina NovaSeq 6000 with paired-end reads. Each sample was aligned to the human genome GRCh38, and analyzed using the 10 × cell ranger 6.0.0, which employed the STAR sequence aligner (Dobin et al., 2013). Cell barcodes were filtered using cellranger 3.0.2, and then downstream analysis was conducted using Seurat (v4.0.1). We integrated two samples using variable genes, and performed downstream analysis. Briefly, low-quality cell data was firstly removed and gene expression matrices were then normalized using the LogNormalized method. Subsequently, we clustered cells using the FindClusters functions and performed non-linear dimensional reduction with the RunUMAP function. Marker genes for each cluster were identified by the "bimod" (Likelihood-ratio test) with default parameters via the FindAllMarkers function in Seurat (v4.0.1). Gene ontology analysis was performed using Enrichr (Kuleshov et al., 2016). Pseudotime trajectories analysis was conducted using the Monocle 2 (v2.18.0) (https://github.com/cole-trapnell-lab/monocle-release).

**Chromatin Immunoprecipitation Sequencing.** The Chromatin Immunoprecipitation (ChIP) assay was executed with the SimpleChIP® Plus Enzymatic Chromatin IP Kit (Magnetic Beads) provided by Cell Signaling Technology (#9005) following the manufacturer's instructions. Briefly, approximately $4 \times 10^6$ cells (equivalent to a 15 cm culture dish) were utilized for each ChIP assay. Initially, the cells were subjected to fixation with 1% formaldehyde for 10 minutes at room temperature. The fixation process was halted by the addition of 2 ml of 10X glycine for 5 minutes. The subsequent steps involved scraping the cells, lysing, digesting, and shearing them by sonication. The lysates were clarified by centrifugation, and 2% of the supernatant was transferred as an 'Input Sample', which could be preserved at -20°C for future use. The remaining lysates supernatant was incubated overnight at 4°C with the immunoprecipitating antibody. This was followed by adding 30 μl of Protein G Magnetic Beads to each IP reaction and incubating them for 2 hours at 4°C with rotation. The beads were then washed, and the chromatin was eluted from the antibody/protein G magnetic beads complex. To all samples, including the 2% input sample from previous steps, 6 μl of 5M NaCl and 2 μl of Proteinase K were added to reverse cross-links. This was followed by a 2-hour incubation at 65°C and DNA purification using Spin Columns. Subsequently, these immuno-enriched

DNA samples were prepared for Next Generation Sequencing (NGS) after constructing a DNA library with the NEBNext® Ultra™ II DNA Library Prep Kit (NEB, USA, Catalog #: E7645L). The qualified libraries were pooled and sequenced on Illumina platforms using a PE150 strategy at Novogene Bioinformatics Technology Co., Ltd (Beijing, China). Finally, the raw reads were subjected to quality control using Fastp and then aligned to the hg19 genome with hisat2. Peak calling was conducted using macs2 and deeptools, and the resultant plots were generated using the IGV software, and the peaks were not normalized by sequencing depth.

**Statistical analysis.** Values are presented as means ± SD. Each figure legend has detailed information explaining the statical test used to show significance. All statistical tests were performed on GraphPad Prism 10, and $P < 0.05$ was considered statistically significant (ns, $P > 0.05$; *$P < 0.05$; **$P < 0.01$; ***$P < 0.001$; ****$P < 0.0001$). Microscopy images were selected form the total quantified images. Microscopy quantified fields were whole fields of well in cell culture plates, and were randomly acquired in immunostained sections.

**Figure 1— figure supplement 1. 2C induced reprogramming of ISL1$^{+}$ cells from hESC-derived CMs.** **(A)** Schematic illustration of optimized protocol to chemically induce and purify CMs differentiated from hESCs. Cells undergoing differentiation from day 0 (D0) to day 8 (D8) were cultured in RPMI1640 basal medium containing B27 without Insulin (RPMI/B-27$^{-INS}$). After 4 days of selection (SD4) in purification medium containing lactate starting from D14, CMs were maintained in RPMI1640 basal medium containing B27 (RPMI/B-27) from maintenance day 0 (MD0) to maintenance day 4 (MD4). Then CMs were used to induce ISL1-expressing cells by treatment with chemicals. **(B)** Expression of ISL1 (green) and CM marker TNNT2 (red) was detected by immunofluorescence staining (left) at SD4, and ISL1$^{+}$ cells and TNNT2$^{+}$ cells were analyzed by flow cytometry analysis (middle) and presented as their percentage (right). Error bars indicate SD. DAPI (4′,6-diamidino-2-phenylindole) staining labeled nuclei as blue. **(C)** Scheme of a large-scale chemical content screening by ISL1 immunofluorescence staining using purified hESC-derived CMs. **(D)** Expression of ISL1 (green) and TNNT2 (red) was shown by immunostaining of CMs treated with various compound combinations for 72 hours. **(E)** Percentage of ISL1$^{+}$ cells after 72-hour treatment of CMs with various compound combinations. Data are shown as mean ± SD. Each treatment condition with 3 technical replicates of values as shown. ns, not significant ($P > 0.05$), ***$P < 0.001$, one-way ANOVA with Dunnett's multiple comparisons test. **(F)** Titration of the A-485 compound on the effects of CM reprogramming in combination of 10μM CHIR99021. The ISL1$^{+}$ and TNNT2$^{+}$ cell percentages were normalized with the negative control DMSO, respectively. Data are shown as mean ± SD (n=3 independent experiments, represented as dots). Two-way ANOVA with Dunnett's multiple comparisons test. ns, not significant ($P > 0.05$), ***$P < 0.001$, ****$P < 0.0001$. **(G)** Changes in cytoplasmic area of CMs following titrated concentrations of A-485 compounds combined with 10μM CHIR99021. Data are shown as mean ± SD (n=3 independent experiments, represented as dots). One-way ANOVA with Dunnett's multiple comparisons test. ****$P < 0.0001$. **(H)** Effects of CM reprogramming induced by the combination of CHIR99021 and I-BET-762. The ISL1$^{+}$ and TNNT2$^{+}$ cell percentages were normalized with the negative control DMSO, respectively. Data are shown as mean ± SD (n=2 independent experiments, represented as dots). Two-way ANOVA with Uncorrected Fisher's LSD. ns, not significant ($P > 0.05$), ****$P < 0.0001$.

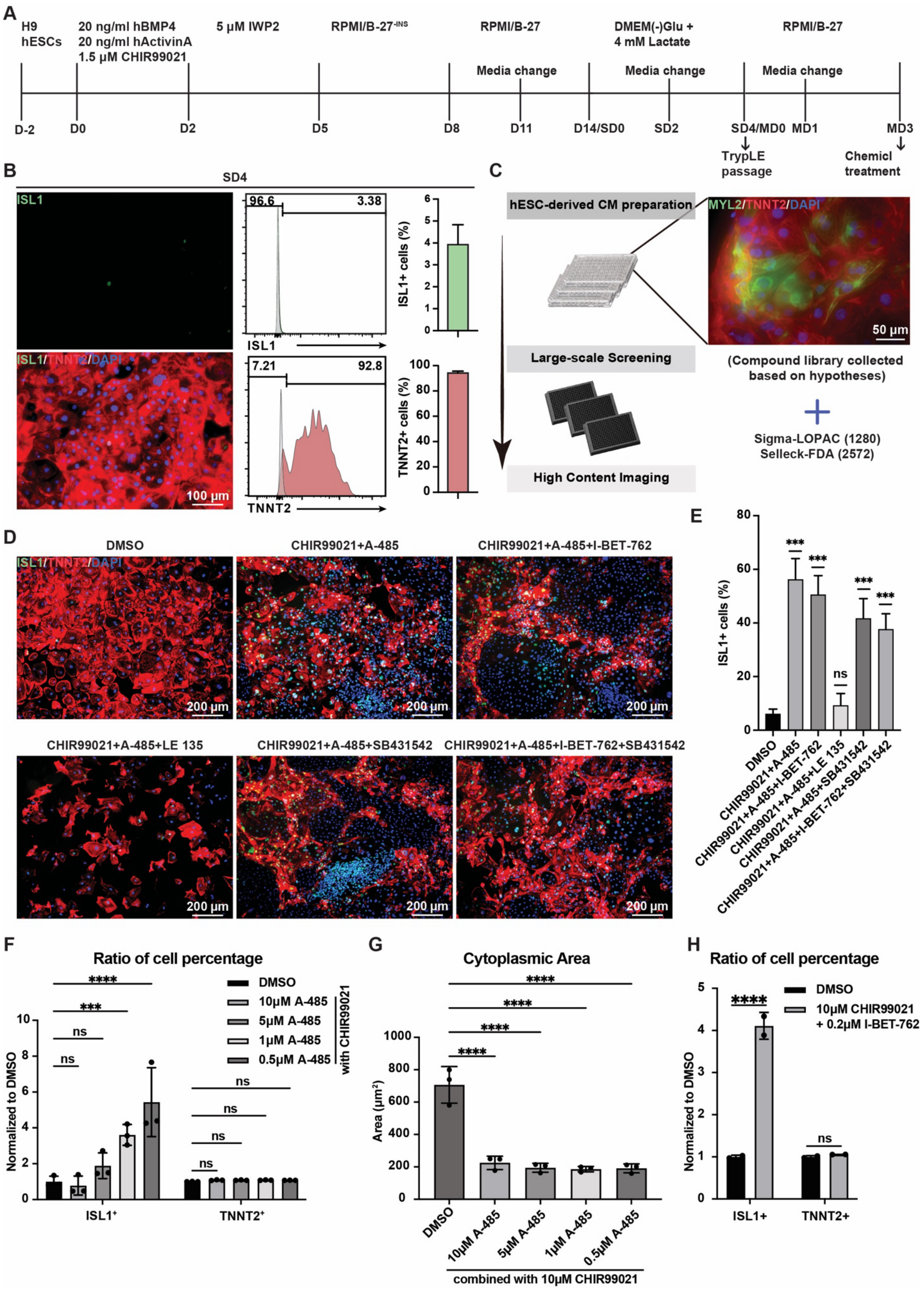
A
H9 hESCs
20 ng/ml hBMP4
20 ng/ml hActivinA
1.5 μM CHIR99021
5 μM IWP2
RPMI/B-27-INS
RPMI/B-27
DMEM(-)Glu +
4 mM Lactate
RPMI/B-27
Media change
Media change
Media change
D-2
D0
D2
D5
D8
D11
D14/SD0
SD2
SD4/MD0
MD1
MD3
TrypLE passage
Chemicl treatment
B
SD4
ISL1
ISL1/TNNT2/DAPI
100 μm
96.6
3.38
ISL1
7.21
92.8
TNNT2
ISL1+ cells (%)
TNNT2+ cells (%)
C
hESC-derived CM preparation
Large-scale Screening
High Content Imaging
MYL2/TNNT2/DAPI
50 μm
(Compound library collected based on hypotheses)
Sigma-LOPAC (1280)
Selleck-FDA (2572)
D
DMSO
CHIR99021+A-485
CHIR99021+A-485+I-BET-762
CHIR99021+A-485+LE 135
CHIR99021+A-485+SB431542
CHIR99021+A-485+I-BET-762+SB431542
ISL1/TNNT2/DAPI
200 μm
E
ISL1+ cells (%)
***
ns
DMSO
CHIR99021+A-485
CHIR99021+A-485+I-BET-762
CHIR99021+A-485+LE 135
CHIR99021+A-485+SB431542
CHIR99021+A-485+I-BET-762+SB431542
F
Ratio of cell percentage
****
***
ns
DMSO
10μM A-485
5μM A-485
1μM A-485
0.5μM A-485
with CHIR99021
Normalized to DMSO
ISL1+
TNNT2+
G
Cytoplasmic Area
****
Area (μm²)
DMSO
10μM A-485
5μM A-485
1μM A-485
0.5μM A-485
combined with 10μM CHIR99021
H
Ratio of cell percentage
****
ns
DMSO
10μM CHIR99021
+ 0.2μM I-BET-762
Normalized to DMSO
ISL1+
TNNT2+

**Figure 1— figure supplement 2. 2C treatment gradually induced de-differentiation of CMs. (A)** Phase contrast images of hESC-derived CMs treated by DMSO (NC) or 2C at indicated time points. **(B)** Gene expression of *ISL1*, *TNNT2*, and *MYL2* at indicated times during 2C treatment. Data are shown as mean ± SD. Gene expression changes at each time point were normalized with the DMSO at the corresponding time point. **(C)** Ratios of $TNNT2^-/ISL1^+$, $TNNT2^+/ISL1^+$, and $TNNT2^+/ISL1^-$ subpopulations at indicated time points after treatment with 2C or DMSO (NC). **(D)** Immunostaining for embryonic cardiogenesis marker genes (MESP1 and NR2F2), pan-cardiac genes (NKX2-5, and GATA4), and ISL1 in DMSO (NC) or 2C-treated CMs for 60 hours. DAPI (4′,6-diamidino-2-phenylindole) staining labeled nuclei as blue. ISL1 (green) is used as a marker of dedifferentiation. **(E)** Immunostaining for ISL1 (green) and CM-specific marker a-Actinin (red) in DMSO (NC) or 2C-treated mature CMs for 60 hours.

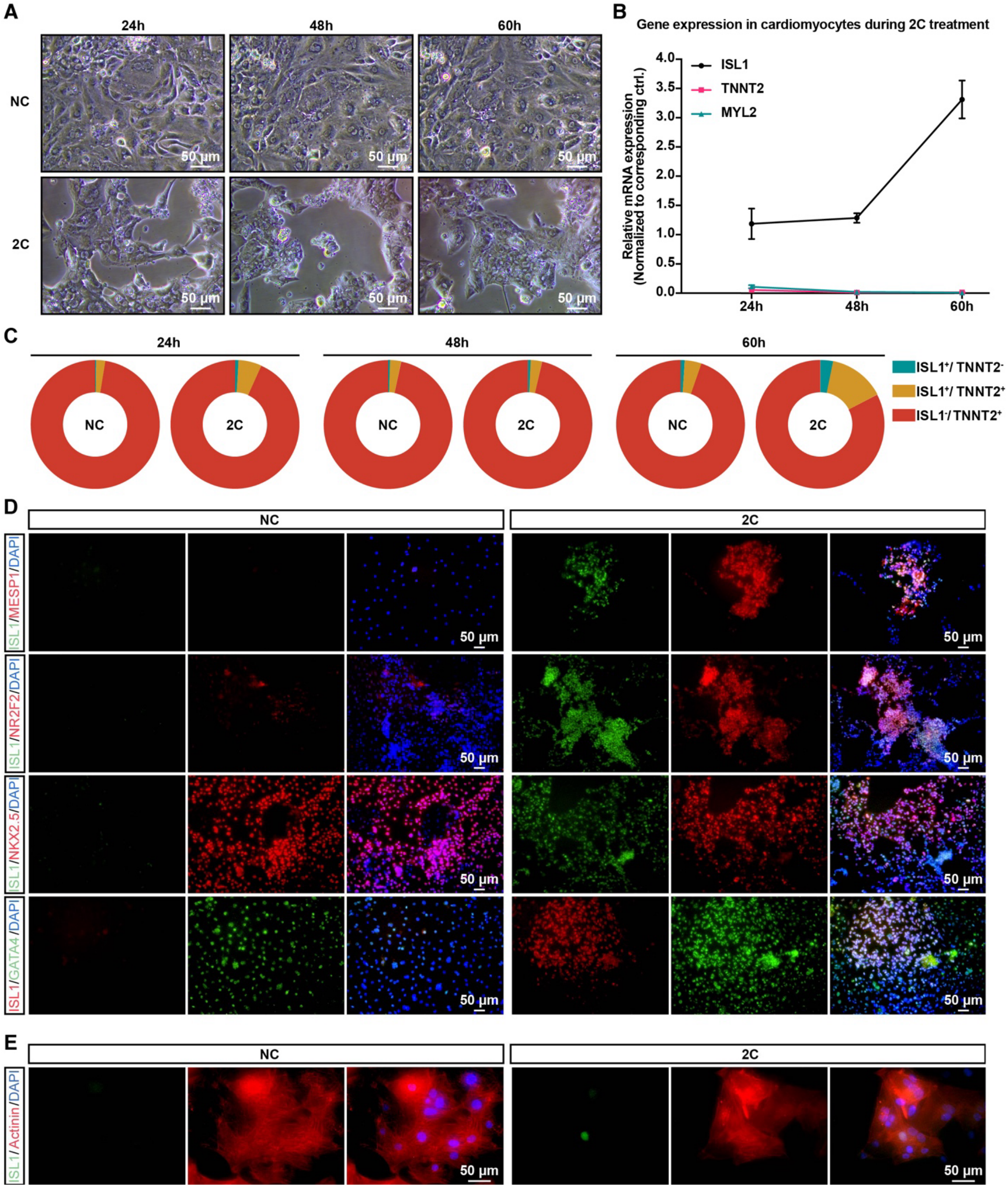
A
24h
48h
60h
NC
2C
50 μm
B
Gene expression in cardiomyocytes during 2C treatment
Relative mRNA expression (Normalized to corresponding ctrl.)
ISL1
TNNT2
MYL2
4.0
3.5
3.0
2.5
2.0
1.5
1.0
0.5
0.0
24h
48h
60h
C
24h
48h
60h
NC
2C
ISL1+/ TNNT2-
ISL1+/ TNNT2+
ISL1-/ TNNT2+
D
NC
2C
ISL1/MESP1/DAPI
ISL1/NR2F2/DAPI
ISL1/NKX2.5/DAPI
ISL1/GATA4/DAPI
50 μm
E
NC
2C
ISL1/Actinin/DAPI
50 μm

270

**Figure 2— figure supplement 1. 2C-induced RCCs represent the differentiation potential ability. (A)** Relative gene expression of embryonic cardiogenesis marker genes (*MESP1*, *ISL1*, *NR2F2*, *FUT4*, and *LEF1*), pan-cardiac genes (*GATA4*, *TBX5*, and *NXK2-5*), and CM marker genes (*MEF2C*, *TNNT2*, *MYL2*, and *MYL7*) in the cells treated by DMSO (NC) or 2C for 60 hours (60h) and subsequently cultured in the absence of 2C for another 3 days (60h+3d). Data are shown as mean ± SD. Multiple unpaired tests. ns, not significant ($P > 0.05$), *$P < 0.05$, **$P < 0.01$, ***$P < 0.001$. **(B)** Flow cytometry analysis of the percentage of CD31⁺ ECs and SMA⁺ SMCs in the cells under corresponding differentiation conditions.

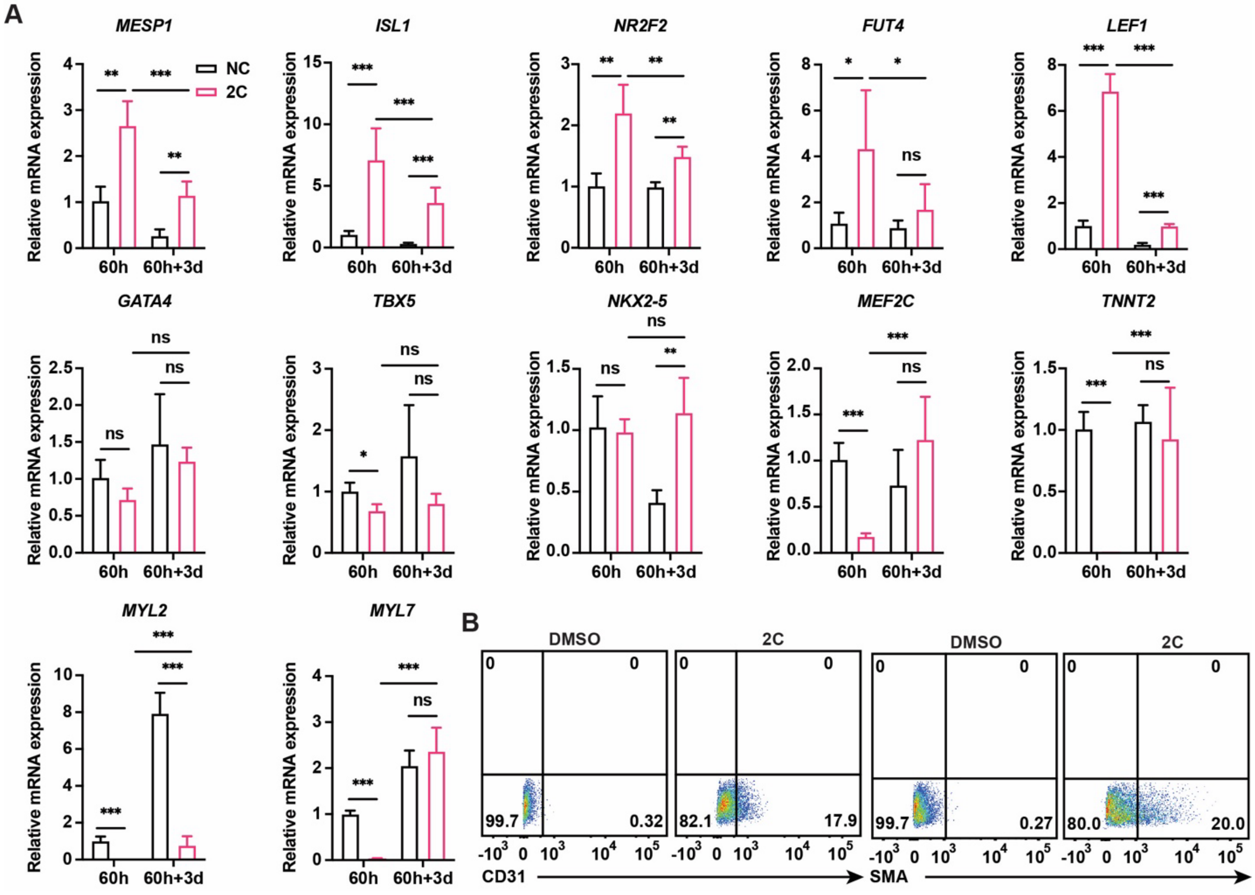

**Figure 3— figure supplement 1. Construction of ISL1mCherry/+ knock-in H9 human ESC line by CRISPR-Cas9. (A)** A diagram showing the strategy to establish H9 hESC line with ISL1-mCherry knock-in reporter. The three single-guide RNAs (Sg1, Sg2, and Sg3) are designed in Exon1 and Intron1 of *ISL1*, respectively. **(B)** Genomic PCR confirmed establishment of heterozygous cell lines with ISL1-mCherry knocked in the targeted ISL1 locus. **(C)** Immunostaining confirmed the expression of ESC marker genes (OCT4, NANOG, and SOX2) and cardiac lineage marker genes (GATA4, NKX2-5, and TBX5) in the cells differentiated from K9 at D0 and D5. **(D)** Relative expression of genes in the cells differentiated from K9 at indicated time points. Error bars indicate SD. **(E)** Immunostaining of CM marker genes (TNNT2, green and MYL2, red) in FACS-sorted K9-derived mCherry-negative CMs. DAPI (4′,6-diamidino-2-phenylindole) staining labeled nuclei as blue.

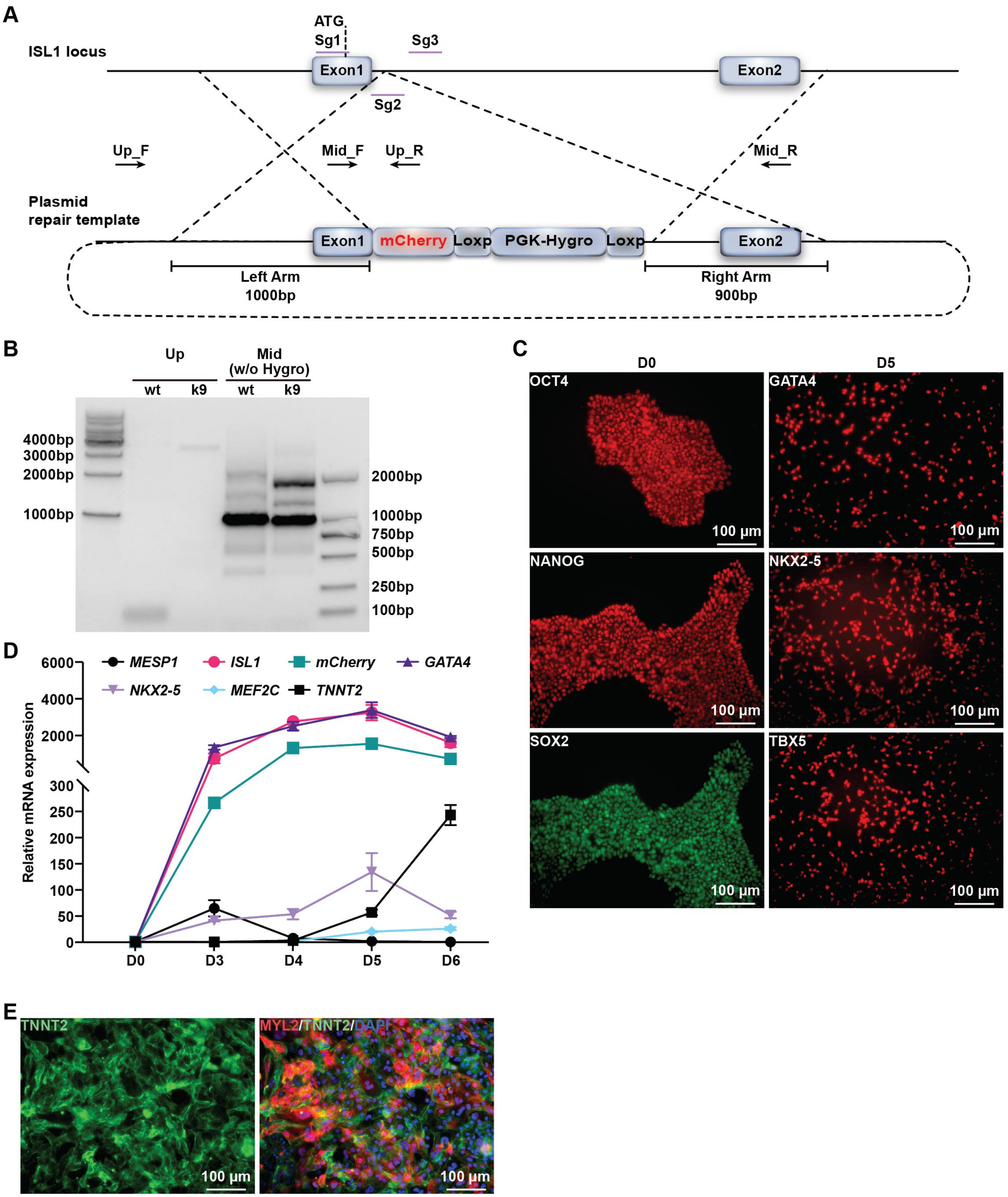
A
ATG
Sg1
Sg3
ISL1 locus
Exon1
Exon2
Sg2
Up_F
Mid_F
Up_R
Mid_R
Plasmid
repair template
Exon1
mCherry
Loxp
PGK-Hygro
Loxp
Exon2
Left Arm
1000bp
Right Arm
900bp
B
Up
Mid
(w/o Hygro)
wt
k9
wt
k9
4000bp
3000bp
2000bp
1000bp
2000bp
1000bp
750bp
500bp
250bp
100bp
C
D0
D5
OCT4
GATA4
NANOG
NKX2-5
SOX2
TBX5
100 μm
D
MESP1
ISL1
mCherry
GATA4
NKX2-5
MEF2C
TNNT2
Relative mRNA expression
6000
4000
2000
300
250
200
150
100
50
0
D0
D3
D4
D5
D6
E
TNNT2
MYL2/TNNT2/DAPI
100 μm

**Figure 3— figure supplement 2. 2C induced K7-derived mCherry-negative CMs into ISL1/mCherry-positive cells. (A)** Immunostaining of ESC marker genes (SOX2, NANOG, OCT4, and SSEA4) and cardiac lineage marker genes (GATA4, NKX2-5, TBX5 and MEF2C) at D0 and D5 of differentiation of $ISL1^{mCherry/+}$ HUES7 knock-in (K7) ESC line. **(B)** Immunostaining showed the co-expression of ISL1 (green) and mCherry (red) in K7-differentiated cells at D6. DAPI (4′,6-diamidino-2-phenylindole) staining labeled nuclei as blue. **(C)** FACS analysis showing the percentage of ISL1/mCherry double positive cells in the cells differentiated from K7 at D6. **(D)** FACS analysis showing the percentage of $mCherry^+$ cells in K7-drived cells cultured in lactate purification media at SD4. **(E)** Flow cytometric analysis showing the percentages of H9 and K7-derived $TNNT2^+$ CMs at SD4. **(F)** FACS analysis showing the percentage of K7-derived mCherry-negative cells at SD4. Error bars indicate SD. **(G)** Immunostaining showed the expression of mCherry and TNNT2 in K7-derived mCherry-negative CMs upon DMSO (NC) or 2C treatment for 60 hours. DAPI (4′,6-diamidino-2-phenylindole) staining labeled nuclei as blue. **(H-I)** Flow cytometry analysis of the percentage of mCherry-positive cells dedifferentiated from K7 hESC KI reporter line derived mCherry-negative CMs. Data are shown as mean ± SD (n=3 independent experiments, represented as dots). **(J)** Relative gene expression of *ISL1*, *mCHERRY*, *LEF1*, *TNNT2* and *MYL2* in K7-derived mCherry-negative CMs treated with DMSO (NC) or 2C for 60 hours, respectively. Data are shown as mean ± SD (n=2 independent experiments with 4 replicates each). Two-way ANOVA with Šidák’s multiple comparisons test. **P < 0.01, ****P < 0.0001.

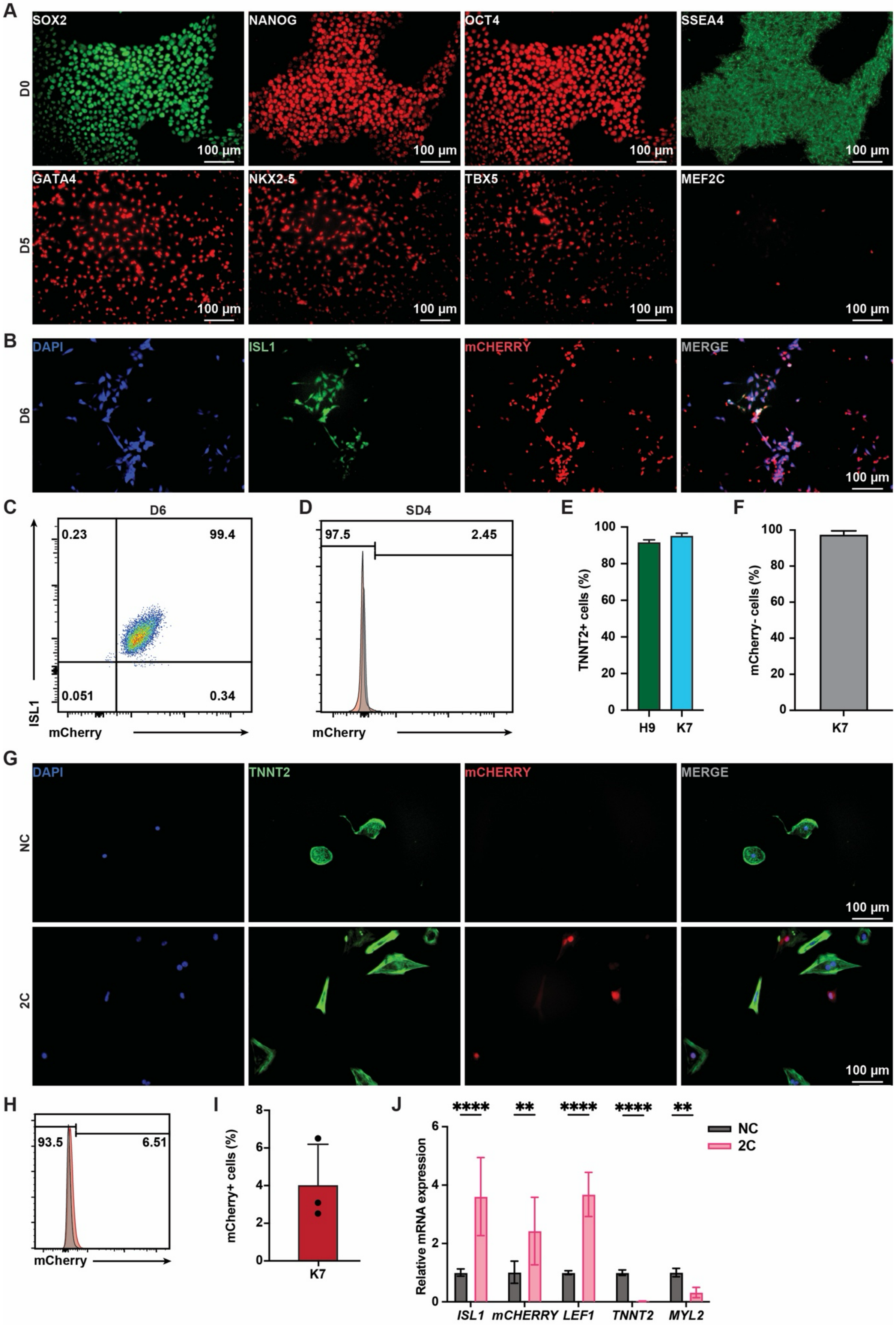
A
SOX2
NANOG
OCT4
SSEA4
D0
GATA4
NKX2-5
TBX5
MEF2C
D5
100 μm
B
DAPI
ISL1
mCHERRY
MERGE
D6
C
D6
0.23
99.4
0.051
0.34
ISL1
mCherry
D
SD4
97.5
2.45
mCherry
E
TNNT2+ cells (%)
H9
K7
F
mCherry- cells (%)
K7
G
DAPI
TNNT2
mCHERRY
MERGE
NC
2C
H
93.5
6.51
mCherry
I
mCherry+ cells (%)
K7
J
NC
2C
Relative mRNA expression
****
**
ISL1
mCHERRY
LEF1
TNNT2
MYL2

**Figure 3— figure supplement 3. 2C induced ISL1 expression in CMs with EGFP labeled by lineage tracing. (A)** Scheme of lineage tracing to label EGFP in CMs. **(B)** Immunostaining showed the expression of EGFP in $TNNT2^+$ CMs infected with or without different titers of virus. DAPI (4′,6-diamidino-2-phenylindole) staining labeled nuclei as blue. **(C)** The number of $TNNT2^+$ cells alive after infection with or without different titers of virus. **(D)** The percentage of $EGFP^+$ cells in $TNNT2^+$ CMs infected with or without different titers of virus.

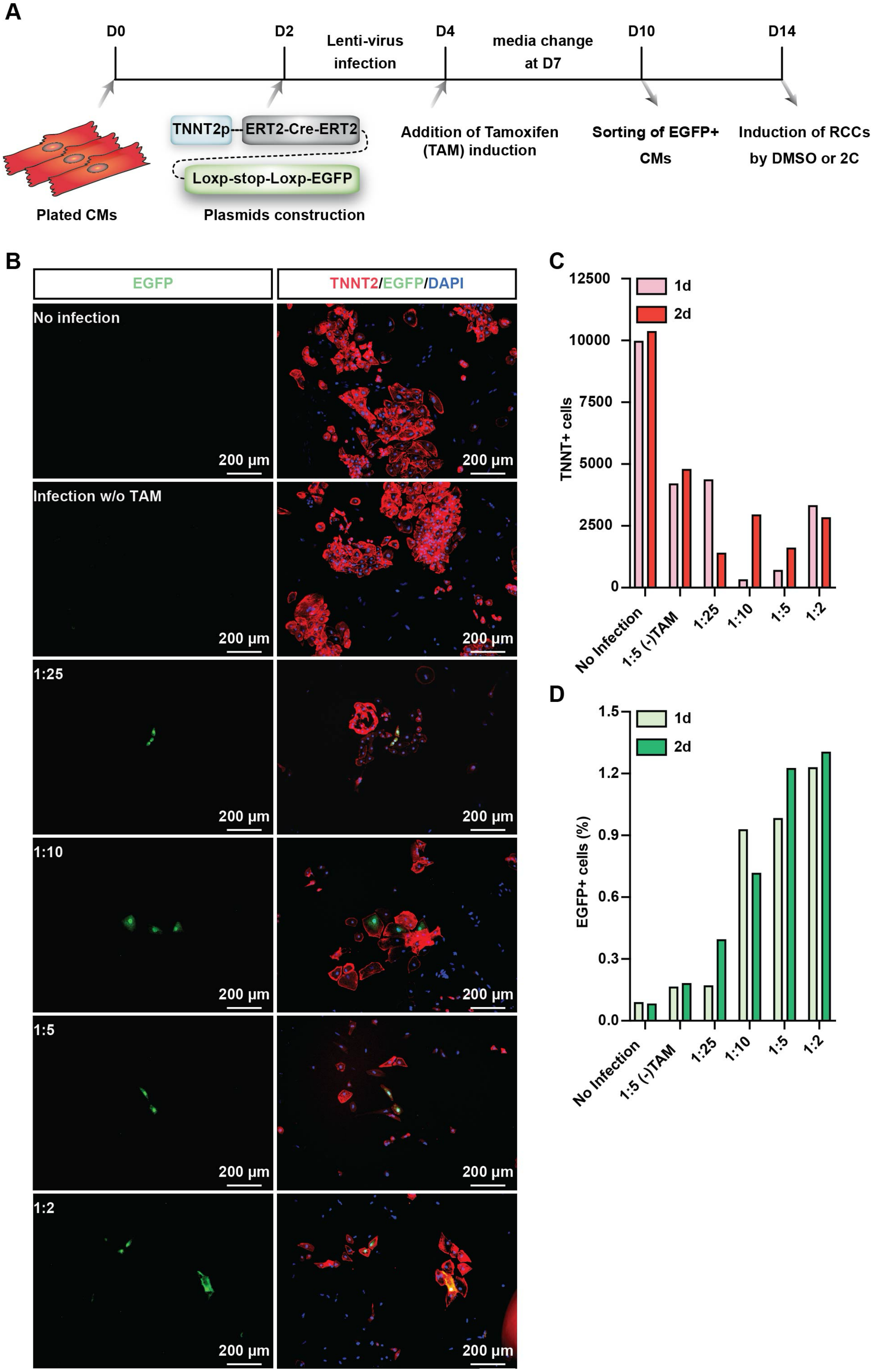
A
D0
D2
Lenti-virus infection
D4
media change at D7
D10
D14
Plated CMs
TNNT2p
ERT2-Cre-ERT2
Loxp-stop-Loxp-EGFP
Plasmids construction
Addition of Tamoxifen (TAM) induction
Sorting of EGFP+ CMs
Induction of RCCs by DMSO or 2C
B
EGFP
TNNT2/EGFP/DAPI
No infection
Infection w/o TAM
1:25
1:10
1:5
1:2
200 μm
C
TNNT+ cells
12500
10000
7500
5000
2500
0
1d
2d
No Infection
1:5 (-)TAM
1:25
1:10
1:5
1:2
D
EGFP+ cells (%)
1.5
1.2
0.9
0.6
0.3
0.0
1d
2d
No Infection
1:5 (-)TAM
1:25
1:10
1:5
1:2

**Figure 4— figure supplement 1. 2C induced expression of ISL1 in neonatal rats CMs *in vitro* and *in vivo*. (A)** Immunostaining of ISL1 (green) and TNNT2 (red) in primary neonatal rats ventricular CMs (NRVCMs) treated with DMSO (NC) or 2C for 60 hours. In 2C-treated cells, cells co-expressing ISL1 and TNNT2 were indicated by yellow arrows. **(B)** Schematic diagram of administration of neonatal SD rats with vehicle (DMSO, NC) or 2C (20 mg/kg CHIR99021 and 10 mg/kg A-485) from the day of birth (P1). **(C-D)** Body weights (**C**) in neonatal rats by administration with DMSO (NC) or 2C at indicated time points and the ratio (**D**) of heart to body weights (HW/BW) under the same condition at day 6. Data are shown as mean ± SD (n=6). Two-way ANOVA with Šidák's multiple comparisons test in (C) and unpaired t test in (D). ns, not significant ($P > 0.05$), **$P < 0.01$. **(E)** Immunofluorescence staining of ISL1 (green) and TNNT2 (red) in cross-sectioned hearts from neonatal rats with the same treatment in (**B**). DAPI (4′,6-diamidino-2-phenylindole) staining labeled nuclei as blue. Ao, aorta. PA, pulmonary artery. LA, left atrial. RA, right atrial. LV, left ventricle. RV, right ventricle.

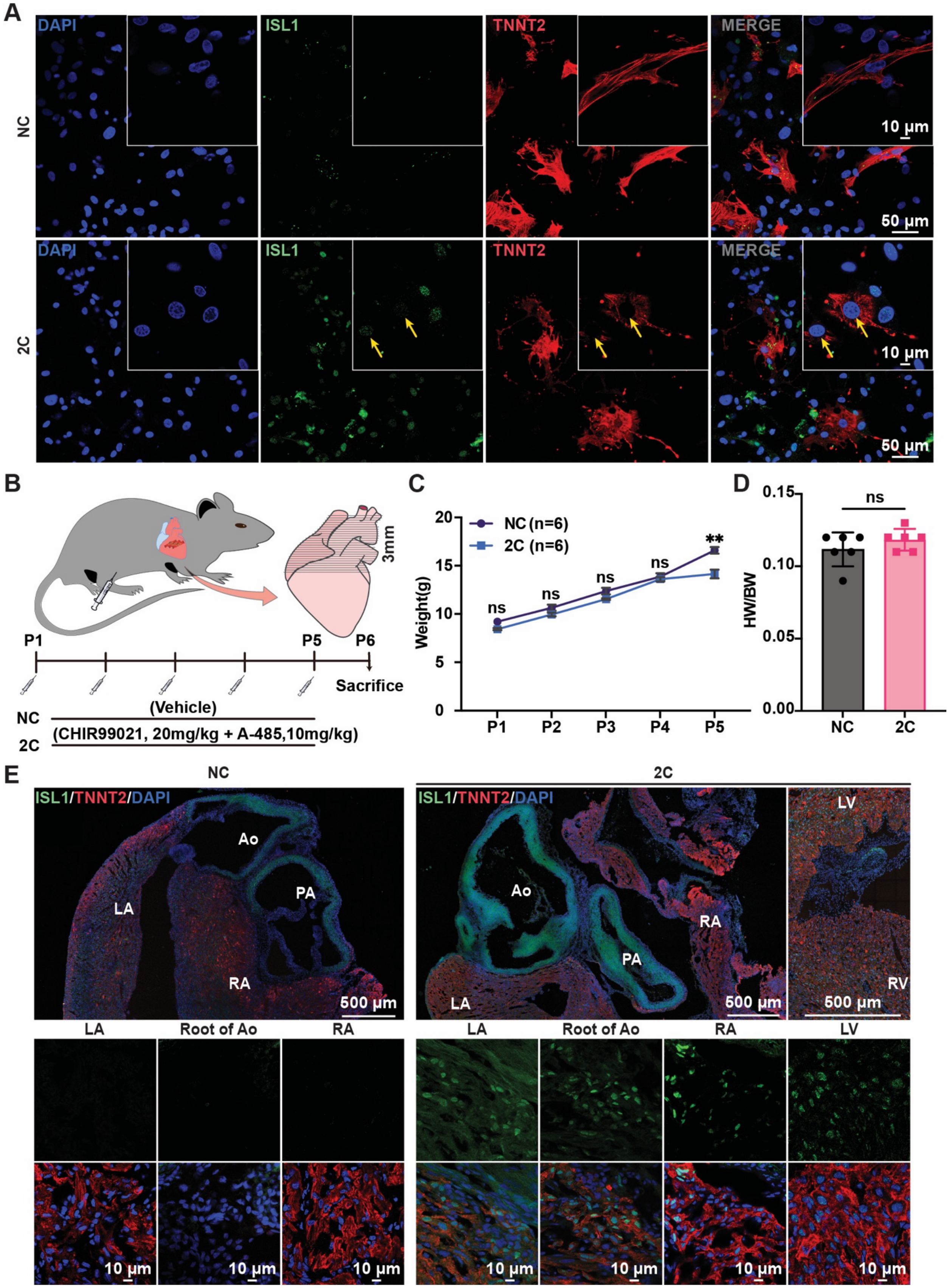

A
DAPI
ISL1
TNNT2
MERGE
NC
2C
10 μm
50 μm
B
3mm
P1
P5
P6
Sacrifice
NC
(Vehicle)
(CHIR99021, 20mg/kg + A-485,10mg/kg)
2C
C
NC (n=6)
2C (n=6)
Weight(g)
ns
**
P1
P2
P3
P4
P5
D
HW/BW
ns
NC
2C
E
NC
2C
ISL1/TNNT2/DAPI
Ao
PA
LA
RA
LV
RV
500 μm
LA
Root of Ao
RA
LV
10 μm

**Figure 4— figure supplement 2. Administration of CHIR99021 or A-485 alone cannot induce ISL1 expression in neonatal rat CMs *in vivo*.** Immunofluorescence staining of ISL1 (green) and TNNT2 (red) in cross-sectioned hearts from neonatal rats with the same treatment in (**Figure 4— figure supplement 1B**). DAPI (4′,6-diamidino-2-phenylindole) staining labeled nuclei as blue. LV, left ventricle. RV, right ventricle.

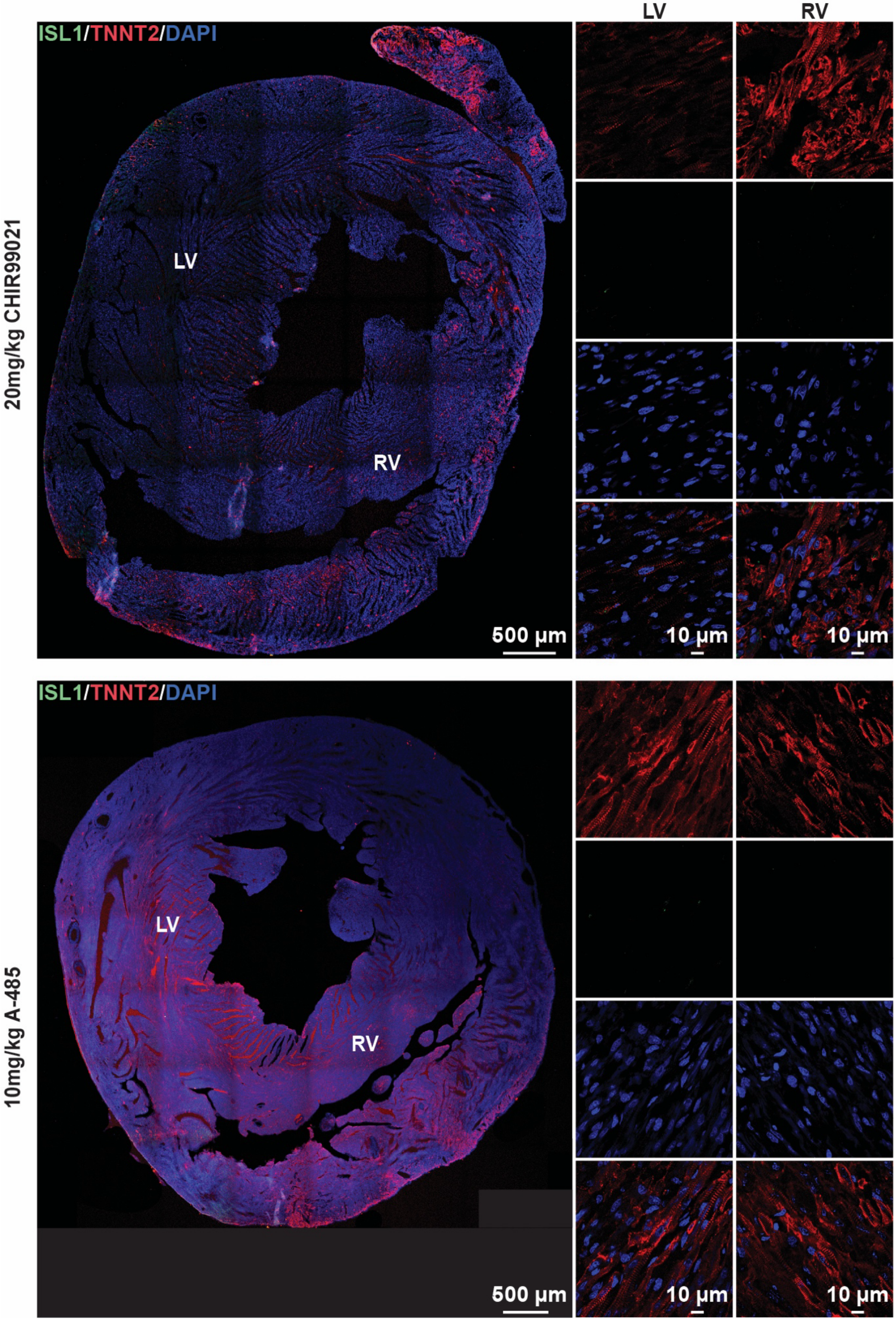

ISL1/TNNT2/DAPI
LV
RV
20mg/kg CHIR99021
LV
RV
500 µm
10 µm
10 µm
ISL1/TNNT2/DAPI
10mg/kg A-485
LV
RV
500 µm
10 µm
10 µm

**Figure 5— figure supplement 1. The individual and cooperative effects from CHIR99021 and A-485 on the induction of H9 hESC-derived CMs into RCCs. (A)** Phase contrast images of hESC-derived CMs treated by DMSO (NC), CHIR99021, A-485, or 2C at indicated time points. **(B-C)** Cytosolic (**B**) and nuclear (**C**) areas of the cells treated by DMSO (NC), CHIR99021, A-485, or 2C at indicated time points. Data are shown as mean ± SD (n=2 independent experiments with 3 replicates each). One-way ANOVA with Dunnett's multiple comparisons test. ns, not significant (P > 0.05), ***P < 0.001. **(D)** Heatmap illustration showing the fold-changes of indicted marker genes' expression by small molecule treatment, which were measured using qRT-PCR. **(E)** Immunostaining showed the expression of ISL1 (green) and TNNT2 (red) in the cells treated by DMSO (NC), CHIR99021, A-485, or 2C for 60 hours (60h) and subsequently cultured in the absence of 2C for another 3 days (60h+3d). DAPI (4′,6-diamidino-2-phenylindole) staining labeled nuclei as blue. **(F)** The ratio of ISL1$^+$ cells in TNNT2$^+$ cells at indicated time points in (**E**). Data are shown as mean ± SD (n=2 independent experiments with 3 replicates each). Two-way ANOVA with Šidák's multiple comparisons test. **(G)** The number of TNNT2$^+$ cells in the cells treated with the same condition in (**E**) at indicated time points. Data are shown as mean ± SD (n=2 independent experiments with 3 replicates each). Multiple unpaired t tests. ns, not significant (P > 0.05), **P < 0.01.

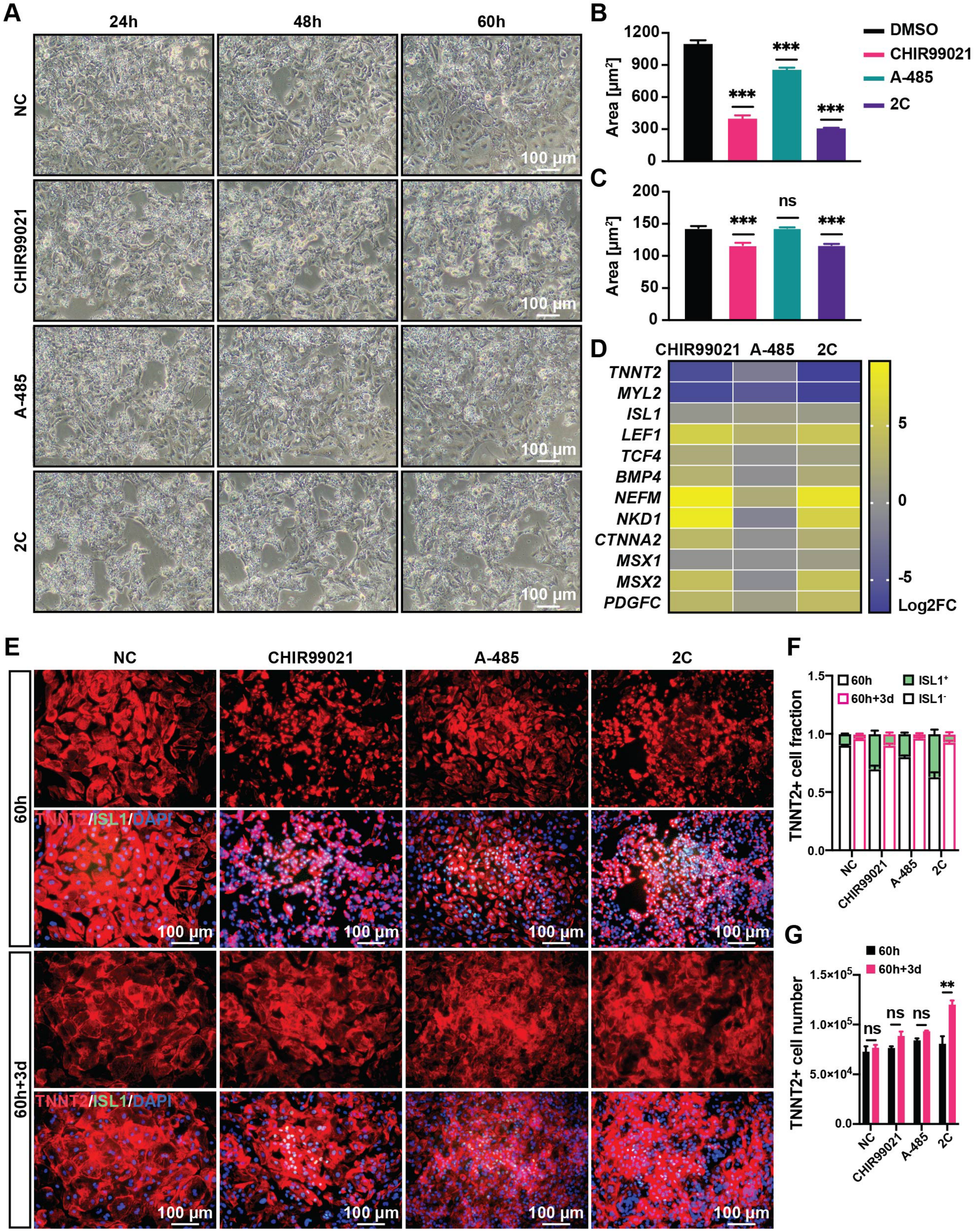
A
24h
48h
60h
NC
CHIR99021
A-485
2C
100 μm
B
Area [μm²]
1200
900
600
300
0
***
DMSO
CHIR99021
A-485
2C
C
200
150
100
50
ns
D
CHIR99021 A-485 2C
TNNT2
MYL2
ISL1
LEF1
TCF4
BMP4
NEFM
NKD1
CTNNA2
MSX1
MSX2
PDGFC
5
0
-5
Log2FC
E
60h
60h+3d
TNNT2/ISL1/DAPI
F
TNNT2+ cell fraction
60h
60h+3d
ISL1⁺
ISL1⁻
1.5
1.0
0.5
0.0
G
TNNT2+ cell number
1.5×10⁵
1.0×10⁵
5.0×10⁴
0.0
**

350

**Figure 7— figure supplement 1. H3K27Ac and H3K9Ac levels regulated by CHIR99021 and A-485 individually or in combination.** Cells induced from H9 hESC-derived CMs by treatment with DMSO (NC), CHIR99021, A-485 or 2C for 60 hours. **(A-C)** H3K27Ac levels in the cells treated with indicated small molecules. Immunostaining (**A**) and quantitative analysis of H3K27Ac levels in the TNNT2$^{+}$ CMs (**B**-**C**). **(D-F)** H3K9Ac levels in the cells treated with indicated small molecules. Immunostaining (**D**) and quantitative analysis of H3K9Ac levels in the TNNT2$^{+}$ CMs (**E-F**). DAPI (4′,6-diamidino-2-phenylindole) staining labeled nuclei as blue. CMs stained by TNNT2 (green). Data are shown as mean ± SD (n=2 independent experiments with 4 replicates each). One-way ANOVA with Dunnett’s multiple comparisons test. ns, not significant ($P > 0.05$), $^{**}P < 0.01$, $^{****}P < 0.0001$.

361 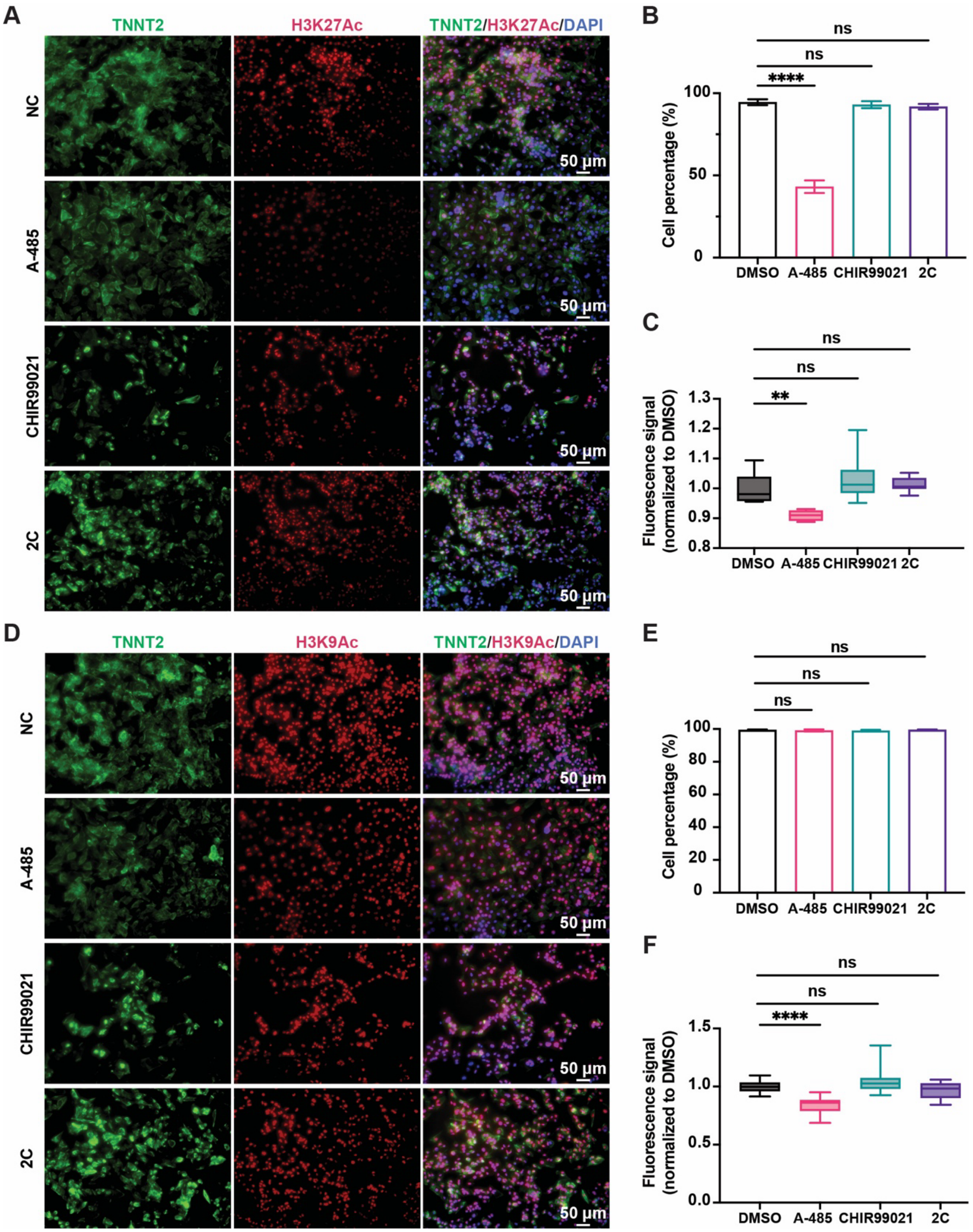
A
TNNT2
H3K27Ac
TNNT2/H3K27Ac/DAPI
NC
A-485
CHIR99021
2C
50 μm
B
ns
ns
****
Cell percentage (%)
100
50
0
DMSO
A-485
CHIR99021
2C
C
ns
ns
**
Fluorescence signal (normalized to DMSO)
1.3
1.2
1.1
1.0
0.9
0.8
DMSO
A-485
CHIR99021
2C
D
TNNT2
H3K9Ac
TNNT2/H3K9Ac/DAPI
NC
A-485
CHIR99021
2C
50 μm
E
ns
ns
ns
Cell percentage (%)
100
80
60
40
20
0
DMSO
A-485
CHIR99021
2C
F
ns
ns
****
Fluorescence signal (normalized to DMSO)
1.5
1.0
0.5
0.0
DMSO
A-485
CHIR99021
2C

**Figure 7— figure supplement 2. Chemical treatment alters the expression of signature genes by regulating H3K27Ac and H3K9Ac in proximity.** Gene ontology (GO) analyses of genes associated with H3K9Ac and H3K27Ac enrichments in the cells treated with DMSO, A-485, CHIR99021, or 2C for 60 hours. **(A-F)** Veen diagram showing the number of genes enriched by H3K9Ac and H3K27Ac after treatment with DMSO, A-485 (**A**-**B**), CHIR99021 (**C**-**D**), or 2C (**E**-**F**). **(G-L)** GO analyses of genes with H3K9Ac and H3K27Ac enrichments observed following treatment with DMSO, A-485 (**G-H**), CHIR99021 (**I-J)**, or 2C (**K-L)**.

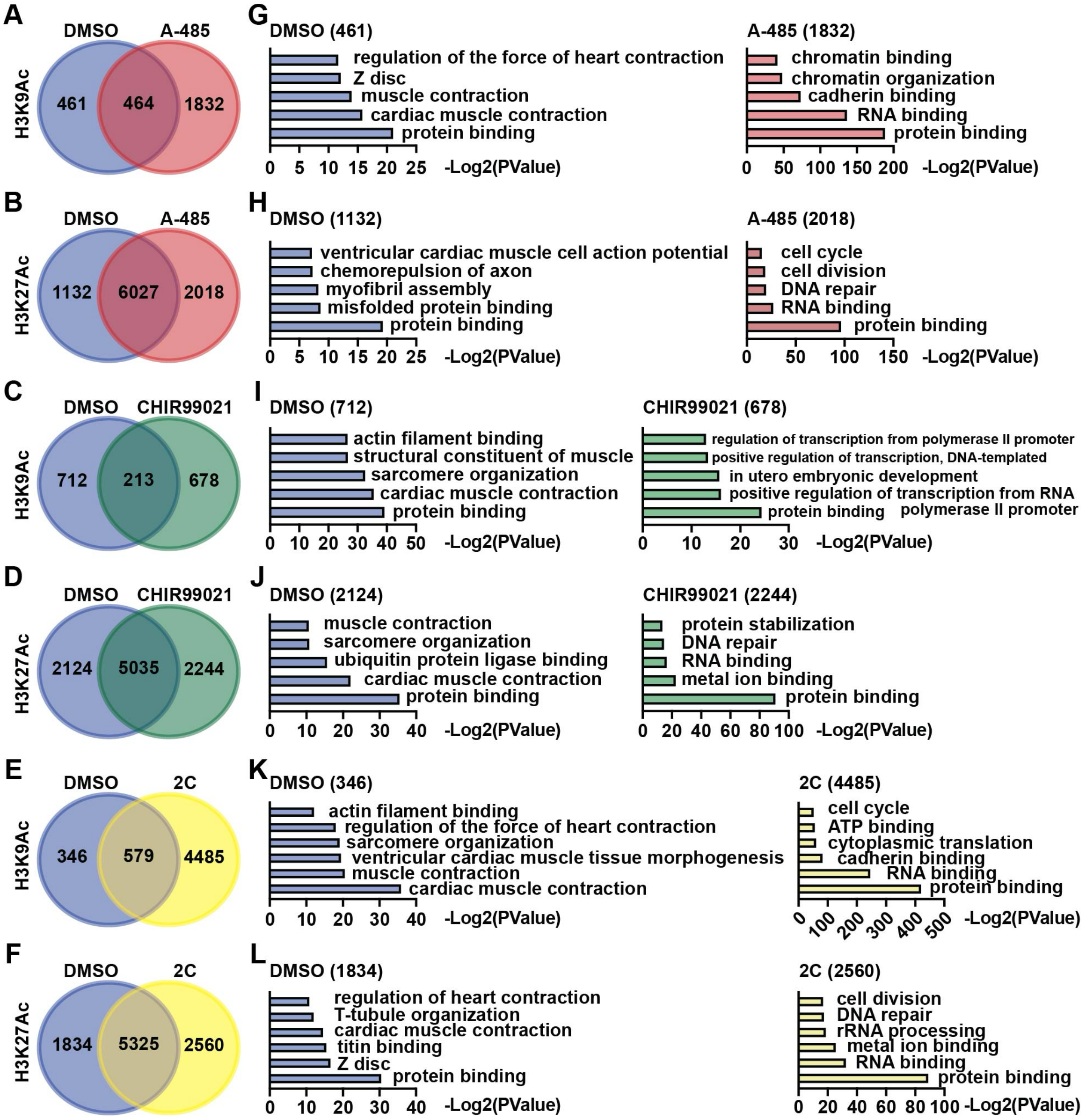

**Figure 7— figure supplement 3. Chemical treatment differentially altered the peaks of H3K9Ac and H3K27Ac on CM and RCC genes. (A)** Peaks of H3K9Ac and H3K27Ac on CM genes in the cells treated with DMSO, A-485, CHIR99021 or 2C, and H3K4me3 peaks on the same genes in the cells treated with DMSO or 2C. **(B-C)** Peaks of H3K9Ac and H3K27Ac on RCC genes in the cells treated with DMSO, A-485, CHIR99021 or 2C, and H3K4me3 peaks on the same genes in the cells treated with DMSO or 2C. The y-axis represents the number of counts.

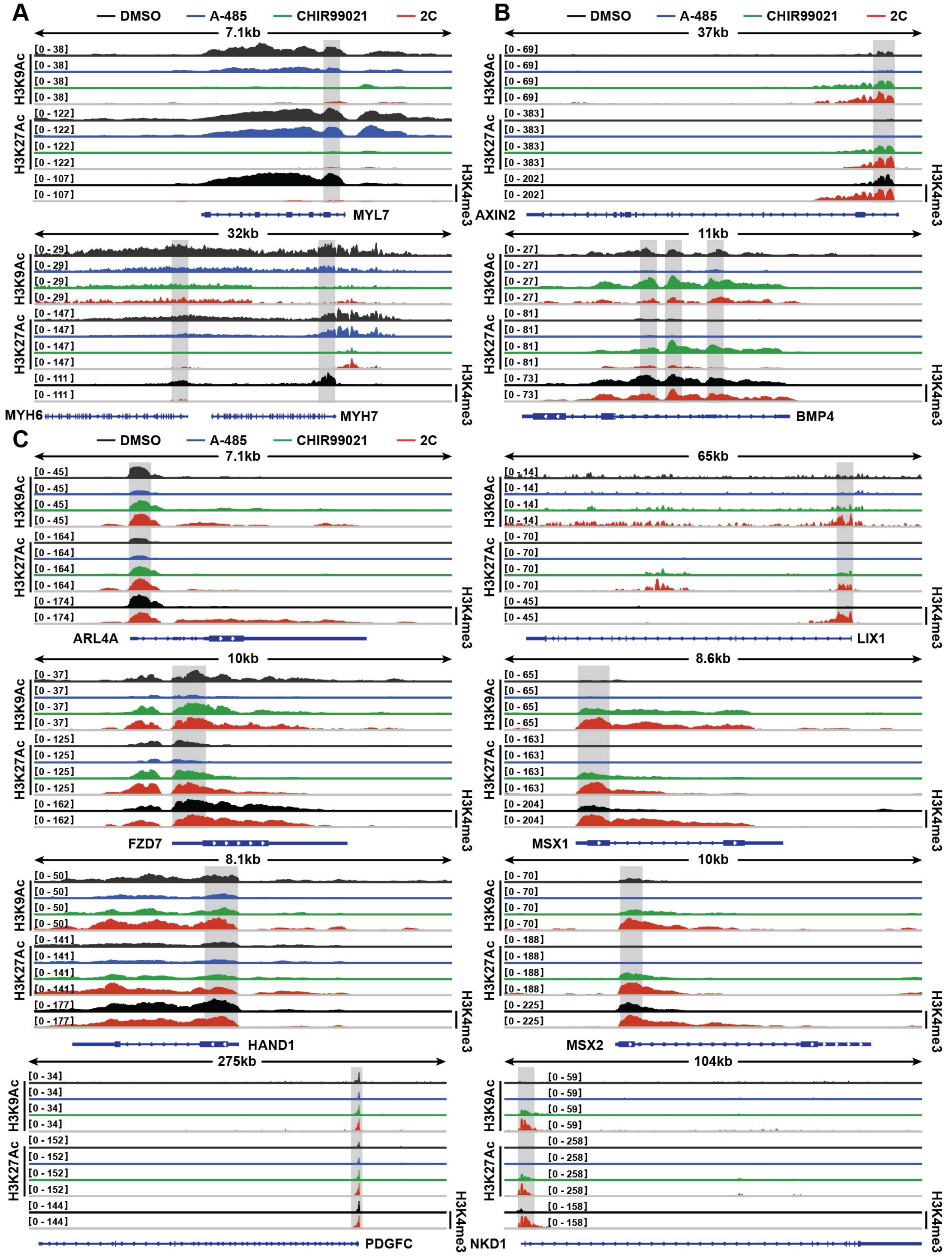

**Tables 1. Compound library collected based on hypotheses.**

| No. | Full Name | Function (s) | Concentration (μM) |
|---|---|---|---|
| 1 | RG108 | DNMTs inhibitor | 0.04 |
| 2 | Decitabine | DNMTs inhibitor | 2 |
| 3 | RSC133 | DNMTs inhibitor | 10 |
| 4 | DZNeP | DNMTs inhibitor | 0.05 |
| 5 | Azacitidine (5AzaC) | DNMTs inhibitor | 2 |
| 6 | Trichostatin A (TSA) | HDACs inhibitor | 0.005 |
| 7 | Sodium butyrate (NaB) | HDACs inhibitor | 250 |
| 8 | Valproic acid sodium salt | HDACs inhibitor | 500 |
| 9 | Vorinostat (SAHA) | HDACs inhibitor | 5 |
| 10 | RGFP966 | HDACs inhibitor | 1 |
| 11 | Romidepsin (FK228) | HDACs inhibitor | 1 |
| 12 | C646 | HAT inhitibor | 2 |
| 13 | BIX01294 | HMTs | 2 |
| 14 | Tranylcypromine HCl (Parnate) | HMTs | 5 |
| 15 | EPZ004777 | HMTs | 5 |
| 16 | SGC 0946 | HMTs | 5 |
| 17 | GSK126 | HMT inhibitor | 10 |
| 18 | GSK-LSD1 2HCl | HMT inhibitor | 10 |
| 19 | ML324 | HDM inhibitor | 5 |
| 20 | GSK J4 HCl | HDM inhibitor | 1 |
| 21 | SB431542 | TGF-b inhibitor | 3 |
| 22 | RepSox | TGF-b inhibitor | 10 |
| 23 | A 83-01 | TGF-b inhibitor | 1 |
| 24 | LY-364947 | TGF-b inhibitor | 5 |
| 25 | LDN-193189 | BMP signaling inhibitor | 0.5 |
| 26 | PD0325901 | MAPK/ERK inhibitors | 1 |
| 27 | Forskolin | PKA activators | 10 |
| 28 | PS48 | PI3K/Akt | 5 |
| 29 | CHIR99021 | Canonical Wnt | 5 |

| | | | |
|---|---|---|---|
| 30 | Apigenin | a potent P450 inhibitor for CYP2C9 | 10 |
| 31 | BIO | GSK-3 Inhibitor | 0.1 |
| 32 | Y27632 2HCl | ROCK inhibitors | 10 |
| 33 | Thiazovivin | ROCK inhibitor | 0.5 |
| 34 | TTNPB | Nuclear Receptor | 1 |
| 35 | AM580 | retinoic acid receptor agonist | 0.01 |
| 36 | Nocodazole (NC) | Cell-cycle inhibitors | 0.1ug/ml |
| 37 | NU6140 (NU) | ATR/CDK inhibitor | 2 |
| 38 | Dasatinib | Src Family Kinase inhibitors | 0.5 |
| 39 | PP1 | Src inhibitor | 5 |
| 40 | Torin1 | Potent and selective mTOR inhibitor | 1 |
| 41 | (R)-(+)-Bay K 8644 | L-type Ca2+-channel blocker | 2 |
| 42 | Kenpaullone | GSK3-beta and CDK inhibitor | 5 |
| 43 | Compound E | Notch signaling suppressor | 0.1 |
| 44 | Vitamin C | Others | 25 ug/ml |
| 45 | L-Ascorbic acid 2-phosphate | stimulate collagen formation | 25 ug/ml |
| 46 | Tretinoin (RA) | a ligand for both the retinoic acid receptor (RAR) and the retinoid X receptor (RXR) | 0.25 |
| 47 | Hh-Ag1.5 | Hedgehog Agonist | 0.5 |
| 48 | Z-VAD-FMK | Cell Death inhibitor | 20 |
| 49 | 3-Methyladenine | Cell Death inhibitor | 5mM |
| 50 | Necrostatin-1 | Cell Death inhibitor | 30 |
| 51 | Liproxstatin-1 | Cell Death inhibitor | 0.2 |
| 52 | Y27632 | Cell Death inhibitor | 10 |
| 53 | Rapamycin | Cell Death inhibitor | 0.1 |
| 54 | IM-54 | Cell Death inhibitor | 10 |
| 55 | diphenyl-benzoquinone (DPQ) | Cell Death inhibitor | 1 |
| 56 | SU5402 | receptor tyrosine kinase inhibitor（VEGFR2、FGFR1、PDGFRβ） | 10 |
| 57 | L-NAME HCl | NO synthase inhibitor | 100 |

| | | | |
|---|---|---|---|
| 58 | JAK inhibitor I | An ATP-competitive inhibitor of Janus protein tyrosine kinases (JAKs). | 1 |
| 59 | SC1 (Pluripotin) | Dual inhibition of ERK1 and Ras GTPase | 1 |
| 60 | PD173074 | FGF receptor inhibition | 0.5 |
| 61 | SU16F | PDGFR-β inhibition | 2 |
| 62 | JNJ10198409 | Dual inhibition of PDGFR-α and PDGFR-β | 0.1 |
| 63 | DAPT | Notch inhibition | 1 |
| 64 | LY-411575 | Notch inhibition | 0.01 |
| 65 | Purmorphamine | Hedgehog activation | 1 |
| 66 | Prostaglandin E2 (PGE2) | PKA activation | 1 |
| 67 | IBMX | PKA activation | 10 |
| 68 | CD437 | RAR activation | 0.1 |
| 69 | Bexarotene | RAR activation | 5 |
| 70 | HX531 | RAR activation | 1 |
| 71 | 9-cis-RA | Dual activation of RAR and RXR | 2 |
| 72 | GW501516 | PPARβ activation | 0.1 |
| 73 | Carbacyclin | PPARβ activation | 10 |
| 74 | IKK 16 | IKK inhibitor | 0.2 |
| 75 | SC-514 | IKK inhibitor | 3 |
| 76 | PF184 | IKK inhibitor | 0.2 |
| 77 | Poly (I:C) | Toll-like receptor 3 (TLR3) activation | 300 ng/ml |
| 78 | Zebularine | DNA methyltransferase inhibition | 100 |
| 79 | UNC0638 | G9a and GLP histone methyltransferase inhibition | 0.5 |
| 80 | Chaetocin | Histone methyltransferase inhibition | 2 |
| 81 | PRT 4165 | Polycomb repressive complex 1 inhibition | 10 |
| 82 | IOX1 | JMJC histone demethylase inhibition | 1 |
| 83 | Tubastatin A | Histone deacetylase inhibition | 0.5 |
| 84 | MS-275 | Histone deacetylase inhibition | 1 |
| 85 | TC-H 106 | Histone deacetylase inhibition | 1 |

| | | | |
|---|---|---|---|
| 86 | MC1568 | Histone deacetylase inhibition | 2 |
| 87 | PCI 34051 | Histone deacetylase inhibition | 0.2 |
| 88 | SIRT1 Inhibitor III | SIRT1 histone deacetylase inhibition | 2 |
| 89 | Salermide | SIRT1/2 histone deacetylase inhibition | 10 |
| 90 | SRT1720 | SIRT1 histone deacetylase activation | 1 |
| 91 | Anacardic acid | Histone acetyltransferase inhibition | 5 |
| 92 | CTPB | P300 histone acetyltransferase activation | 5 |
| 93 | JQ1 | BET bromodomain inhibition | 0.2 |
| 94 | I-BET-762 | BET bromodomain inhibition | 0.2 |
| 95 | OAC1 | Epigenetic modulation | 10 |
| 96 | OAC2 | Epigenetic modulation | 5 |
| 97 | N-oxaloylglycine | Prolyl 4-hydroxylase inhibition | 1 |
| 98 | Quercetin | mitochondrial ATPase and phosphodiesterase inhibition | 1 |
| 99 | 2-Deoxy-D-glucose | Glycolysis inhibition | 5000 |
| 100 | Fasudil (HA-1077) HCl | ROCK inhibition | 2 |
| 101 | Pyrintegin | Integrin signaling activation | 3 |
| 102 | Eosin Y Disodium Trihydrate (AMI-5) | Histone arginine methyltransferase inhibition | 5 |
| 103 | CD1530 | Potent and selective RARγ agonist | 0.1 |
| 104 | DY131 | A selective agonist at ERRβ and ERRγ | 10 |
| 105 | DLPC | NR5A2 agonist | 50 |
| 106 | Ch55 | RAR-a/b activator | 1 |
| 107 | SMER28 | regulator of autophagy | 10 |
| 108 | AS8351 | KDM5B inhibitor | 1 |
| 109 | Resveratrol | SIRT1 histone deacetylase activation | 5 |
| 110 | Pifithrin-α (PFTα) | P53 inhibition | 5 |
| 111 | Pifithrin-μ | P53 inhibition | 5 |
| 112 | 17β-Estradiol | ESR activator | 10 |
| 113 | Torkinib (PP242) | mTOR inhibitor | 1 |

| | | | |
|---|---|---|---|
| 114 | BMS-189453 (RAi) | Synthetic retinoid and RARβ agonist; also RARαand RARγ antagonist | 1 |
| 115 | LY294002 | PI3K inhibitor | 1 |
| 116 | LOE 908 hydrochloride | a broad spectrum cation channel blocker | 5 |
| 117 | A23187, free acid | Calcium ionophore | 1 |
| 118 | Phorbol 12-myristate 13-acetate (PMA) | Protein kinase C activator | 0.1 |
| 119 | SNAP | A stable analog of endogenous S-nitroso compounds | 100 |
| 120 | SR 202 | Selective PPARγ antagonist | 5 |
| 121 | LE 135 | Retinoic acid antagonist | 5 |
| 122 | NKH 477 | Water-soluble analog of forskolin | 5 |
| 123 | PAC-1 | Activator of procaspase-3; pro-apoptotic | 5 |
| 124 | GSK 4716 | Selective agonist of ERRβ and ERRγ | 5 |
| 125 | ML 228 | HIF pathway activator | 5 |
| 126 | Acetylcysteine (NAC) | Glutathione (GSH) precursor and cell permeable antioxidant | 5 |
| 127 | Pentamidine isethionate (PTM) | inhibits constitutive nitric oxide synthase in the brain and acts as a NMDA glutamate receptor antagonist | 5 |
| 128 | DMOG | α-KG antagonistand HIF prolylhydroxylase inhibitor | 5 |
| 129 | Roscovitine (Seliciclib,CYC202) | a potent, selective inhibitor of CDK | 10 |
| 130 | Aloisine A RP107 (CAS 496864-16-5) | An inhibitor of CDK1, CDK2, CDK5, GSK-3 alpha, and JNK | 0.1 |
| 131 | RPI-1 | RET Receptor Tyrosine Kinase Inhibitor | 10 |
| 132 | GW3965 HCl | Active non-steroidal agonist for the liver X receptor (LXR) | 2 |
| 133 | T0901317 | LXR agonist | 10 |
| 134 | 24(S)-Hydroxycholesterol (EPM-1) | endogenous agonist for LXR | 1 |
| 135 | Pregnenolone-16α-carbonitrile (PCN) | PXR (pregnane X receptor) activator | 10 |
| 136 | SR 12813 | PXR agonist | 2 |

| | | | |
|---|---|---|---|
| 137 | Cytosporone B | Naturally occurring NR4A1 agonist | 1 |
| 138 | Ciglitazone | Selective agonist at PPARγ | 10 |
| 139 | SR 1664 | High affinity PPARγ ligand; blocks Cdk5-dependent PPARγ phosphorylation | 1 |
| 140 | Genistein | PPARγligand,estrogen receptor ligand and EGFR inhibitor | 2 |
| 141 | Pirfenidone | Antifibrotic agent | 10 |
| 142 | Nintedanib (BIBF 1120) | Inhibits multiple tyrosine kinases | 1 |
| 143 | Rosiglitazone | Potent and selective PPARγ agonist | 10 |
| 144 | Pioglitazone | Selective PPARγ agonist | 1 |
| 145 | Imatinib (STI571) | Inhibitor of tyrosine kinases of the TGFβand PDGF pathways | 10 |
| 146 | SIS3 | Selective Smad3 inhibitor | 5 |
| 147 | Calpeptin | calpain inhibitor | 0.1 |
| 148 | CITCO | Constitutive androstane receptor agonist | 2 |
| 149 | Bumetanide (EPM-3) | MET | 5 |
| 150 | Estradiol valerate (EPM-2) | MET | 5 |
| 151 | CAS 313981-82-7 (EPM-11) | MET | 5 |
| 152 | CAS 912791-92-5 (EPM-13) | MET | 5 |
| 153 | CAS 890825-02-2 (EPM-15 ) | MET | 5 |
| 154 | Lanosterol (EPM-4) | Cholesterol precursor sterol | 5 |
| 155 | Tamoxifen | Estrogen receptor partial antagonist | 10 |
| 156 | Oxindole I | A potent, selective inhibitor of VEGF | 10 |
| 157 | Methacycline HCl | MET | 5 |
| 158 | AUTEN67 | MTMR inhibitor | 10 |
| 159 | SF51 | calcium channel atagonist | 20 |
| 160 | NC043 | USP30 inhibitor | 2 |
| 161 | EPI743 | CoQ10 analogue | 1 |
| 162 | Urolithin A | Mitophagy inducer | 50 |
| 163 | Doxycycline hyclate | an inhibitor of matrix metallo-proteinases (MMP) | 1 |

| | | | |
|---|---|---|---|
| 164 | LiCL | inhibits the replication of type 1 and type 2 Herpes | 2 |
| 165 | Nicotinamide | PARP-1 inhibitor | 10 |
| 166 | sphingosine-1-phosphate | A lipid second messenger that binds to S1P1 and S1P3 receptors | 0.5 |
| 167 | Ponasterone A | derivates of ecdysone, a kind of insect hormone | 1 |
| 168 | Blebbistain | myosin II ATPase inhibitor | 2 |
| 169 | RO4929097 | γ secretase inhibitor, Notch inhibitor | 10 |
| 170 | Oleoyl-L-a-lysophosphatidic acidic sodium salt | a proliferative and anti-apoptotic factor, signaling for PI3K-mediated regulation of cell activity. | 5 |
| 171 | D-Fructose 1,6-bisphosphate trisodium salt | An allosteric activator of enzymes | 5 |
| 172 | EPZ015666 | Prmt5 inhibitor | 5 |
| 173 | MI-2 | MLL2(MLL) inhibitor | 10 |
| 174 | SGI-1027 | Dnmt3A/B inhibitor | 100 |
| 175 | MK-5108 (VX-689) | AURORA_A inhibitor | 1 |
| 176 | AZD1152-HQPA | AURORA_B inhibitor | 1 |
| 177 | 1400W dihydrochloride | iNOS inhibitor | 100 |
| 178 | FH1(BRD-K4477) | hepatocyte maturation activator | 25 |
| 179 | FPH1 (BRD-6125) | hepatocyte maturation activator | 25 |
| 180 | QNZ | NF-kB inhibitor | 5 |
| 181 | Wortmannin | PI3K inhibitor | 1 |
| 182 | Dexamethasone | Dexamethasone | 0.1 |
| 183 | Luteolin | TNF-α, IL-6, NF-Kb, AP-1 inhibitor | 7.5 |
| 184 | WL5A5 | MST inhibitor | 1 |
| 185 | Clomiphene citrate | estrogen agonist | 2 |
| 186 | Niclosamide | anthelmintic and potential antineoplastic activity | 0.1 |
| 187 | Kinetin | a geroprotector and a cytokinin | 20 |
| 188 | Fluphenazine | a phenothiazine and antipsychotic agent | 10 |
| 189 | PFI 3 | an azabicycloalkane | 2 |
| 190 | LY2090314 | GSK-3α/β inhibitor | 5 |

| | | | |
|---|---|---|---|
| 191 | CP2 | KDM4 inhibitor | 5 |
| 192 | Dorsomorphin 2HCl (Compound C) | a potent and selective inhibitor of AMPK | 2 |
| 193 | XAV939 | Tankyrase1/2 inhibitor | 5 |
| 194 | IWP2 | Porcn mediated Wnt palmitoylation | 5 |
| 195 | A-485 | p300/CBP selective catalytic inhibior | 10 |
| 196 | ascorbic acid | increases the active iron (Fe2+) required for the TET | 10 |
| 197 | CHIR-98014 | GSK-3α/β inhibitor | 10 |
| 198 | adenosine | metabolite | 1 |
| 199 | 4-Hydroxyquinoline | metabolite | 100 |
| 200 | Fumaric acid | metabolite | 100 |
| 201 | SAG | Smo receptor agonist | 1 |
| 202 | TTFA | complex II inhibitior | 5 |
| 203 | ISRIB | inhibition of ISR | 5nM |
| 204 | Eriodictyol 7-O-glucoside | unknown | 10 |
| 205 | vitamin K1 | photosynthesis | 10 |
| 206 | WP1066 | apoptosis | 5 |
| 207 | ciclopirox | iron chelator | 2 |
| 208 | BIRB | P38 | 10 |
| 209 | Bergapten | cell replication | 5 |
| 210 | BIA 2-093 | antiepileptic | 5 |
| 211 | Ketanserin tartrate | 5-HT2A | 5 |
| 212 | Doxazosin Mesylate | apoptosis | 5 |
| 213 | Elesclomol (STA-4783) | oxidative stress inducer | 5 |
| 214 | Flupirtine maleate | analgesic | 1ug/ml |
| 215 | Ropinirole HCl | anti-oxidant | 5 |
| 216 | T0070907 | PPARγ | 20 |
| 217 | Loteprednol etabonate | antiinflammation | 5 |
| 218 | Epoxomicin | proteasome inhibitor | 5 |
| 219 | Oleuropein | proteasome activator | 5 |
| 220 | MG132 | proteasome inhibitor | 5 |

| | | | |
|---|---|---|---|
| 221 | Crizotinib | ALK inhibitor | 5 |
| 222 | Ceritinib | ALK inhibitor | 5 |
| 223 | Brigatinib | ALK inhibitor | 5 |
| 224 | Lapatinib | RTK inhibitor | 5 |
| 225 | Vemurafenib | MAPK inhibitor | 10 |
| 226 | ABT-263 | anti-apoptotic protein inhibitor | 5 |
| 227 | WM-1119 | KAT6A inhibitor | 1 |
| 228 | Dabrafenib | MAPK inhibitor | 1 |
| 229 | Trametinib | MAPK inhibitor | 1 |
| 230 | Timapiprant | DP2 antagonist | 5 |
| 231 | Ridaforolimus (deforolimus) | mTOR inhibitor | 5 |
| 232 | GW441756 | Tropomyosin-related kinase A (TrkA) inhibitor | 5 |
| 233 | ZLN005 | PGC-1α transcriptional activator | 10 |
| 234 | Tracheloside | decrease the activity of alkaline phosphatase | 10 |
| 235 | Periplocin | activation of Src/ERK,PI3K/Akt | 10 |

**Movie 1. Contracted CMs at SD4.** hESC-derived CMs at day 4 (SD4) after lactate selection.

**Movie 2. Contracted CMs at 60h+3d-2C.** hESC-derived CMs treated by2C for 60 hours (60h) and subsequently cultured in the absence of 2C for another 3 days (60h+3d).
